# Calibration of AlphaGenome Variant Impact and SpliceAI2 scores as PP3/BP4 evidence for missense, non-coding, and synonymous variants

**DOI:** 10.64898/2026.10.21.760715

**Authors:** Ali Babazade, Yarkın Dolaş, Tarik Duzenli, Ozan Vural, Ceren Furtana, Mehmet Ali Ergun

**Affiliations:** Gazi University, Faculty of Medicine, Department of Medical Genetics, Ankara, Türkiye

**Author notes:** These authors contributed equally (Ali Babazade, Yarkın Dolaş). Corresponding author. Address: Emniyet District, Mevlana Boulevard, No:29, 06560, Yenimahalle/Ankara, Türkiye.

**Keywords:** AlphaGenome, SpliceAI2, in silico, variant classification, PP3/BP4

## Abstract

Computational predictions can serve as graded evidence in variant classification after calibration. We calibrated the AlphaGenome Variant Impact score (AVI) for missense variants and for six non-coding and synonymous classes, and SpliceAI2 for the six classes. For missense variants, AVI reached evidence between moderate and strong in both directions but did not outperform its component AlphaMissense. AVI reached strong pathogenic evidence in the three classes adjacent to exons. It reached moderate evidence, the highest estimable level, in proximal introns and at other synonymous sites, and supporting evidence, at an unstable threshold, in deep introns. SpliceAI2 reached the same levels, except one level lower at near-canonical positions and none in deep introns. No reached level failed validation on later ClinVar classifications, although 14 of 30 benign levels were not tested. With population variants as controls, both tools reached supporting and moderate pathogenic evidence in all six classes, at stricter thresholds near exons and at other synonymous sites, and at lower thresholds in proximal and deep introns. ClinVar likely benign classifications changed about 60-fold in number at 20 nucleotides from the exon without a change in measured splicing. AVI, a genome-wide score, did not differ detectably in discrimination from the splicing-specific SpliceAI2 in four classes; SpliceAI2 discriminated better in two classes and in a splicing assay, and was more sensitive at 99% specificity in four classes. Both scores provide calibrated evidence for non-coding and synonymous variants, but thresholds depend on whether ClinVar benign classifications or population variants are used for comparison.

## Introduction

The American College of Medical Genetics and Genomics and Association for Molecular Pathology (ACMG/AMP) framework allows computational predictions to count as evidence for the pathogenicity of a variant (PP3) or against it (BP4).^1^ In its original form this evidence had a single, supporting strength, and the choice of tools and cutoffs was left to the individual laboratories. A subsequent Bayesian interpretation of the framework assigned likelihood ratios (odds of pathogenicity) to the different evidence strengths.^2^ Pejaver et al. and later Bergquist et al. applied this approach to calibrate missense predictors against ClinVar classifications, allowing score intervals to be assigned supporting, moderate, or strong evidence strengths.^3,4^ This approach has since been extended to in-frame insertions and deletions^5^ and, with population controls, to variants in 5′ cis-regulatory regions.^6^

For splicing predictions, by contrast, the current recommendations rest on a different outcome. The Clinical Genome Resource (ClinGen) Sequence Variant Interpretation (SVI) Splicing Subgroup derived SpliceAI^7^ cutoffs from variants with splicing-assay results, with altered splicing as the outcome and the same cutoffs at all positions outside the canonical dinucleotides.^8^ This approach does not depend on clinical classifications, but altered splicing is, as the authors note, an imperfect surrogate for pathogenicity: a variant that changes splicing in an assay does not necessarily cause disease. Moreover, whether a score carries the same weight of evidence for pathogenicity at a near-canonical splice position, in the polypyrimidine tract, and deep in an intron has not been established. At the same time, calibration against clinical classifications, as used for missense variants, has its own limitations in these regions. Known pathogenic non-coding variants are few, and benign classifications are concentrated at the positions that laboratories routinely report.^9^ In addition, some intronic and synonymous variants classified as benign have been shown to alter splicing.^9^

Two recently released models score variants outside the coding sequence. AlphaGenome^10^ predicts regulatory and splicing effects from sequence. Its developers also provide the AlphaGenome Variant Impact score (AVI) for every single-nucleotide variant (SNV): a supervised combination of features, including AlphaMissense,^11^ trained on allele-frequency labels from the Genome Aggregation Database (gnomAD) on a subset of chromosomes.^12^ SpliceAI2, by contrast, is a splicing model trained on RNA sequencing data from ten mammalian species, with precomputed scores for all SNVs in protein-coding genes.^13^ A recent preprint compared AVI with splicing predictors and measured evidence levels for intronic and untranslated variants together.^14^ Neither model has yet been calibrated separately for each position class, and no thresholds have been proposed for PP3/BP4.

Here, we calibrated AVI for missense variants and for six position-defined non-coding and synonymous classes, and SpliceAI2 for the same six classes with the same pipeline. The missense set contained 5,310 pathogenic or likely pathogenic and 3,108 benign or likely benign variants, and the six non-coding and synonymous classes contained 3,277 pathogenic or likely pathogenic variants. Because the reliability of benign classifications in introns was itself in question, every class was also calibrated with rare population variants as controls. Finally, we examined the calibrated thresholds against a saturation splicing assay of 310,405 substitutions across approximately 600 exons.^15^ The results show that, with ClinVar benign classifications as the benign class, both tools can provide strong pathogenic evidence, or evidence between moderate and strong, in the classes adjacent to exons. However, the thresholds depend on whether ClinVar benign classifications or population controls are used.

## Material and methods

### Study design

AVI was calibrated for missense, non-coding, and synonymous variants, whereas SpliceAI2 was calibrated for the non-coding and synonymous classes (Figure 1). Each calibration comprised a main analysis, which gives the proposed thresholds; robustness analyses, which test how the thresholds change; and descriptive analyses. All data analyzed in this study were public and aggregated, and no new data were collected from human participants.

**Figure 1.**
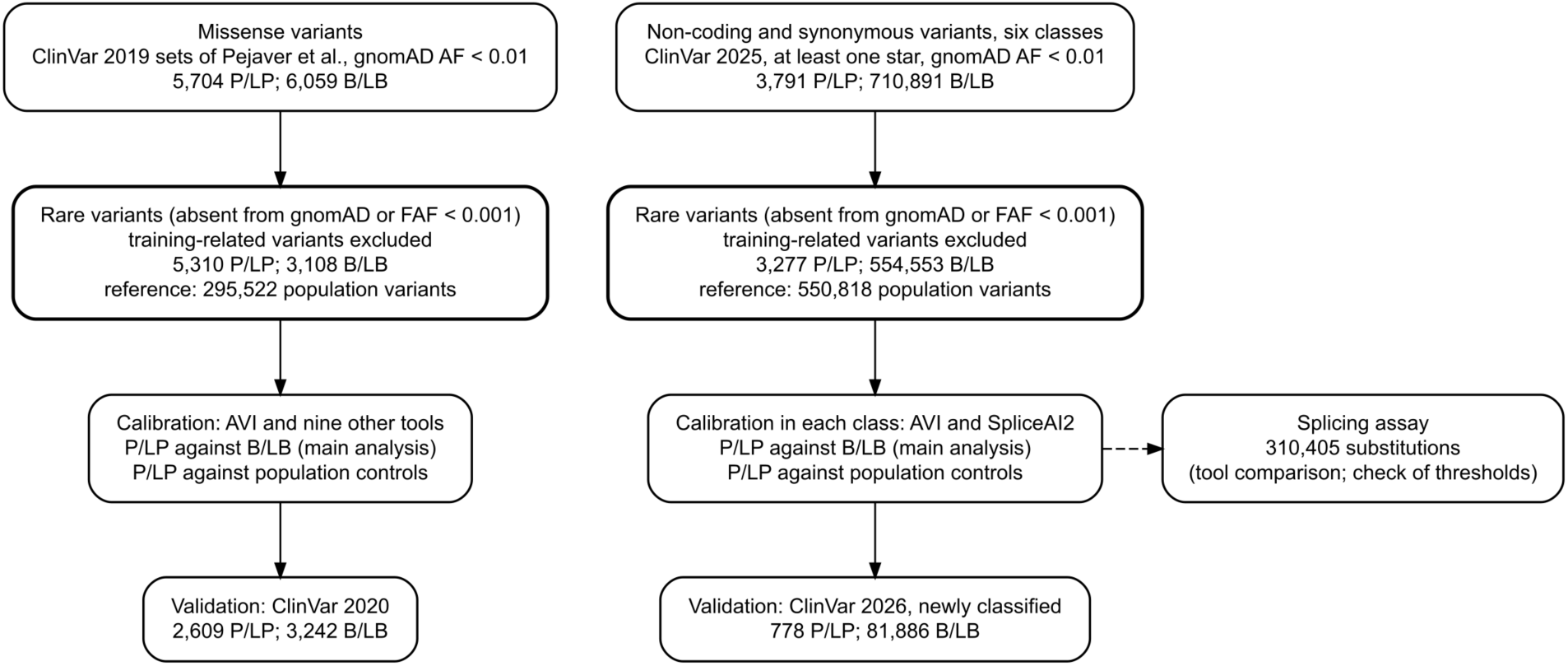
Study design and variant flow. Left, missense variants; right, non-coding and synonymous variants. For both variant groups, pathogenic or likely pathogenic (P/LP) variants were compared with benign or likely benign (B/LB) variants (main analysis) and, separately, with rare population variants as controls. The thresholds were then tested on later classifications. Reference, rare population variants that the method uses to set the window size and that serve as controls. The splicing assay was used to compare the tools and to check the thresholds. AVI, AlphaGenome Variant Impact score; AF, allele frequency; gnomAD, Genome Aggregation Database; FAF, filtering allele frequency.

### Scores

AVI scores were obtained from the genome-wide file of SNVs released with the AlphaGenome Atlas.^12^ Thresholds were calculated on the raw scores and are reported on the PHRED scale of the AVI release (supplemental methods). SpliceAI2 scores were taken from the precomputed scores released with SpliceAI2 (version 2.0);^13^ we used the summary score, the largest of the four predicted changes in splice-site use (acceptor or donor gain or loss), and, for variants with scores on both strands, the higher value.

### Calibration and evidence levels

We used the local posterior probability method of Pejaver et al.,^3,4^ in our own implementation, which closely reproduced their published thresholds (supplemental methods). For each candidate threshold, a local positive likelihood ratio (LR) is estimated in the smallest score window around it that contains at least 100 labeled variants and at least 3% of a reference set of population variants. In the Bayesian adaptation of the ACMG/AMP framework,^2^ each evidence level worth *n* points requires a minimum LR of *c^n/8^*. Following Pejaver et al., *c* was set to 1,124, derived from a prior probability of pathogenicity of 4.41% for rare missense variants.^3^ The resulting minimum LRs for pathogenic evidence were 2.406 (supporting, 1 point), 5.790 (moderate, 2 points), 13.93 (3 points), and 33.53 (strong, 4 points), and the maximum LRs for benign evidence were their reciprocals: 0.416, 0.173, 0.072, and 0.030.

The 3-point evidence level falls between moderate and strong evidence in the points-based framework.^16^ Following recent calibrations,^4,5^ we refer to this level as +3 and to the corresponding benign evidence level as −3. Reported thresholds are one-sided 95% bootstrap bounds; therefore, an evidence level was counted as reached only if it was reached in at least 95% of bootstrap replicates, which makes the thresholds conservative. In addition, a level was not counted if its threshold was reached by no more than one labeled variant of its direction. We applied the same criteria to all classes. Posterior probabilities also depend on the prior probability of each class; we therefore estimated a range of priors for each class and report the corresponding probabilities in the Results. We also recalculated the likelihood-ratio requirements for priors of 0.5%, 1%, 2%, and 10% (supplemental methods).

### Missense variants

We used the calibration (ClinVar 2019), reference (gnomAD v2), and validation (ClinVar 2020) sets published by Pejaver et al. and Bergquist et al.^3,4^ To minimize potential overlap with AVI training data, the primary set excluded variants at positions on AVI training chromosomes where gnomAD v4.1 genomes reported a variant, variants in four genes that the AVI developers used to select the final model, and variants at positions included in a benchmark set.^17^ We refer to these exclusions as the training-related exclusion. We further restricted the analysis to rare variants (the rarity rule: absent from gnomAD v4.1 genomes or with a filtering allele frequency below 0.001).^18^ The resulting set contained 5,310 pathogenic or likely pathogenic and 3,108 benign or likely benign variants.

Published thresholds for other tools had been derived using less restricted variant sets. We therefore recalibrated nine other tools using the same variants and procedure. These included the seven tools recommended by ClinGen for PP3/BP4 evidence (BayesDel,^19^ MutPred2,^20^ REVEL,^21^ VEST4,^22^ AlphaMissense,^11^ ESM1b,^23^ and VARITY_R^24^),^3,4^ as well as CADD v1.6^25^ and v1.7.^26^ For the primary set, we performed 10,000 bootstrap replicates by resampling variants, following Pejaver et al.,^3^ so that the thresholds are comparable with published calibrations. AVI may perform better on variants it saw in training. To check this, we compared its performance on its training chromosomes with that on the eight chromosomes used neither for training nor for model selection. We ran the same comparison for AlphaMissense, BayesDel, and CADD v1.6, whose training did not separate these chromosomes from the others (supplemental methods). We also repeated the AVI calibration in robustness analyses that each changed one choice (supplemental methods).

### Non-coding and synonymous variants

Calibration labels were germline ClinVar classifications with at least one review star in the 15 June 2025 release, which the AVI developers also used for evaluation.^12,27^ We included SNVs in genes with at least one pathogenic or likely pathogenic classification in this release, with an allele frequency below 0.01 in gnomAD v4.1 genomes, and applied the same training-related exclusion and rarity rule as for missense variants.

Variants were annotated on Matched Annotation from NCBI and EMBL-EBI (MANE) Select transcripts (GENCODE v46).^28^ Each intronic and synonymous variant was assigned to one of seven classes: near-canonical splice positions (donor +3 to +6 and acceptor −3), polypyrimidine tract (acceptor −4 to −20), donor flank (+7 to +20), proximal intronic (21–100 nucleotides from the exon), deep intronic (more than 100 nucleotides from the exon), exonic splice region (first base or last three bases of an exon; synonymous variants only), and other synonymous variants. The splice-region positions follow Walker et al.,^8^ and the 100-nucleotide boundary follows a common definition of deep intronic variants.^29^ The six classes other than the donor flank were calibrated; together they contained 3,277 pathogenic or likely pathogenic and 554,553 benign or likely benign variants. Promoter variants (up to 1,000 nucleotides upstream of the transcription start site) and untranslated region variants were included as exploratory classes, and canonical splice-site positions (±1 and ±2) were excluded. Position +5 accounts for approximately half of pathogenic near-canonical variants, so we also performed a calibration excluding this position (Table S1).

The evidence levels that could be estimated depended on the number of labeled variants. A class was calibrated only if it had at least 400 labeled variants, with at least 100 in the smaller label group. Levels above moderate were estimated only for classes with at least 1,600 labeled variants and at least 300 in the smaller group.

The reference set, which sets the window size in the calibration and from which the population controls were drawn, consisted of quality-filtered gnomAD v4.1 genome variants of the same classes and genes on chromosomes not used for AVI training, excluding variants present in ClinVar (sampling in the supplemental methods).

Calibration used 2,000 gene-cluster bootstrap replicates, which resample genes rather than variants, so that variants of the same gene are not treated as independent. Gene-cluster and variant-level resampling were compared for missense variants and for the non-coding and synonymous classes (Tables S1 and S2a).

### Validation

For each evidence level reached during calibration, we calculated the LR of the corresponding score interval using variants with later ClinVar classifications. Uncertainty was estimated using the 5th and 95th percentiles of the Jeffreys posterior distribution (90% credible interval). For missense variants, we used the ClinVar 2020 validation set of the published sets. For the non-coding and synonymous classes, we selected variants classified as pathogenic, likely pathogenic, benign, or likely benign in the ClinVar release of 28 September 2026 but not in the 2025 release, with the same training-related exclusion and rarity rule. This included both newly submitted variants and previously reported variants that had subsequently been reclassified, such as those initially classified as variants of uncertain significance. Because previously submitted variants may have been re-evaluated for reasons related to their prediction scores, we repeated the validation using only first-time ClinVar submissions, that is, variants absent from the 2025 release. Neither validation set excluded variants whose classifications may have incorporated computational predictions.

Verdicts were assigned in the following order, with *L* the required LR. A level was not tested if its score interval was empty or if, given the number of variants in the interval, fewer than five benign variants would be expected there at an LR of exactly *L*. A level failed if the 95th percentile was below *L*. A level was validated if the 5th percentile was at least *L*, probably validated if the point estimate was at least *L*, and unclear if the point estimate was below *L* but the 95th percentile was at least *L*. The same criteria were applied in reverse for benign evidence. A level that failed would not have been reported as reached; the other verdicts did not change the reported levels. The benign variants in the validation set were selected using the same criteria as those in the calibration set; therefore, this analysis could not determine whether the thresholds depended on how benign variants were selected.

### Population variants as controls

The main analysis used ClinVar benign classifications. To assess robustness, we repeated the calibration using rare population variants from the reference set as controls, following the approach of Villani et al. for 5′ regulatory variants.^6^ Because population controls may include undiscovered pathogenic variants, the estimated likelihood ratios are biased toward 1 and cannot exceed the reciprocal of the pathogenic fraction among the controls (the likelihood-ratio cap). We report uncorrected results and considered a pathogenic level assessable only when its required likelihood ratio was no greater than half this maximum, using the estimated upper limit of the pathogenic fraction of each class or, for missense variants, 4.41%; where a level was not reached but its requirement exceeded half this maximum, it was reported as not assessable rather than as not reached (supplemental methods). Corrected analyses assuming pathogenic fractions of 0.5%–10% are given for the non-coding and synonymous classes in Table S1; for missense variants, the correction assumed 4.41%.

Computational evidence is used mainly for rare variants, because common variants can be classified by their population frequency. We therefore restricted both ClinVar benign variants and population controls to rare variants. AVI was trained on allele-frequency labels. For such scores, Bergquist et al. recommended calibration within the most stringent frequency group, as in our main analysis, or separate calibrations across frequency groups.^4^ We therefore also repeated the calibration using higher-frequency population variants as controls (filtering allele frequency ≥0.001 in at least one ancestry group and global allele frequency <0.01). These variants are less likely to be pathogenic, but they fall in the frequency range that AVI was trained to treat as neutral, so this analysis is not independent of AVI. As relatively few higher-frequency variants were located near exons, we compared their thresholds with those obtained from sets of the same size, drawn at random 20 times each from ClinVar benign classifications and from rare population variants.

### Classifications made before SpliceAI

ClinVar classifications of non-coding and synonymous variants may have used SpliceAI predictions since SpliceAI was published in January 2019.^7^ We therefore divided the calibration set of each class into two groups: variants that already had the same classification, with at least one review star, in the ClinVar release of 2 December 2018, and all other variants. Most variants in the second group were classified or reclassified later. We compared likelihood ratios under the main thresholds and the areas under the receiver operating characteristic curve (AUCs) of SpliceAI2 and AVI between the two groups. In addition, we repeated the calibration using only the later classifications and evaluated every level reached against the earlier classifications with the validation rules (supplemental methods).

### Robustness analyses

For the non-coding and synonymous classes, we repeated the calibration in robustness analyses that each changed one choice, including the training-related and rarity exclusions (with matched random removal), the label set, the reference window (1% instead of 3%), exclusion of donor position +5, the boundary between proximal and deep introns (200 and 500 nucleotides), the bootstrap unit, and restriction of both ClinVar classes to variants absent from gnomAD v4.1 genomes. The full list is given in the supplemental methods.

### Comparison of tools

We evaluated further tools on a fixed subset of variants chosen without using any score (supplemental methods): CADD v1.7,^26^ phyloP based on 241 mammals as a conservation-only baseline,^30^ SpliceAI computed with a maximum distance of ±4,999 nucleotides and as precomputed scores obtained through the Ensembl VEP REST API,^7^ and SpliceTransformer;^31^ the original SpliceAI was also calibrated with the same pipeline. AVI and SpliceAI2 were compared on variants scored by both tools using the AUC and sensitivity at 99% specificity, with either ClinVar benign variants or population variants as the negative class and 2,000 gene-cluster bootstrap replicates; these comparisons were descriptive, without adjustment for multiple testing. To assess incremental value, we measured the gain in AUC when one score was added to the other in logistic regression models. To explore the contribution of AVI beyond splicing prediction, we counted variants with an AVI score at or above the pathogenic supporting threshold of their class and a SpliceAI2 score below 0.05, overall and among classifications made before SpliceAI. We also calibrated the four intronic classes together, to assess whether pooled thresholds carry comparable evidence across classes (pooled thresholds are not proposed), and calculated the likelihood ratio of all original SpliceAI scores ≥0.2 taken together, the cutoff of Walker et al.^8^ For missense variants, AVI was compared with AlphaMissense and REVEL in the same way, and we calculated rank correlations of AVI with AlphaMissense, CADD v1.7, and REVEL (supplemental methods).

### Splicing assay

We used the per-substitution results of the saturation minigene assay of Quarantani et al.,^15^ which measured exon inclusion for 310,405 SNVs across approximately 600 exons (Supplementary Table 4 of version 1 of the preprint). A substitution was classified as disruptive if its percent spliced in was at most half that of the wild type and the assay’s significance criterion was met. We calculated AUCs for each position class outside the canonical splice-site dinucleotides (near-canonical splice positions, polypyrimidine tract, donor flank, intronic positions 21 or more nucleotides from the exon, and exonic positions) and compared AVI with SpliceAI2 using 2,000 exon-cluster bootstrap replicates. Each substitution in a calibrated class was placed in the evidence interval that contains its score, separately for each tool. For each interval, we calculated a likelihood ratio for splice disruption, the share of disruptive substitutions falling in the interval divided by the same share for non-disruptive substitutions, following Walker et al.^8^ We also compared the share of disruptive substitutions at each intronic position with the number of ClinVar classifications at that position.

### Benign classifications, population controls, and prior probability

We calculated the share of rare population variants falling within each evidence interval. For proximal and deep intronic variants, we compared the shares of ClinVar pathogenic, ClinVar benign, and population variants at or above the population-control thresholds within bins of distance from the exon and within the same genes; for deep intronic variants, we also measured how concentrated the variants were in genes and recalculated AUCs without the genes with the most pathogenic variants. The prior probability of pathogenicity of each class was bounded below by a floor, the share of rare population variants already classified as pathogenic. It was bounded above by a ceiling, an upper confidence limit that assumes that undiscovered pathogenic variants score like known ones. We also estimated it with a mixture-proportion method (DistCurve^32^) using five other prediction scores (CADD v1.7, phyloP, SpliceAI with the ±4,999-nucleotide window and as precomputed scores, and SpliceTransformer; supplemental methods).

### Software

All analyses were performed in Python 3.13; SpliceAI and SpliceTransformer scores were computed in Python 3.11 and 3.10, respectively.

## Results

### AVI reached +3 and −3 evidence for missense variants

In the primary set, AVI reached supporting, moderate, and +3 pathogenic evidence and supporting, moderate, and −3 benign evidence, but did not reach strong evidence in either direction (Table 1; Figure 2; Table S2a). All six levels were validated using ClinVar 2020 variants (Table S2b). The same levels were reached in the published sets and with gene-cluster bootstrapping, and supporting and moderate evidence were reached in all robustness analyses (Tables S2a and S2c). AVI showed no sign of better performance on variants it may have seen in training: the difference in performance between its training chromosomes and the chromosomes used neither for training nor for model selection was similar in size and direction to that of three tools whose training did not separate these chromosomes (supplemental methods). Additionally, AVI assigned pathogenic evidence to 14% and benign evidence to 52% of rare population missense variants (Table S3). An AVI PHRED score of 20, a cutoff often used with PHRED-scaled scores, fell within the benign supporting interval (19.60 to 21.98), so an AVI score of 20 is not a pathogenicity threshold for missense variants.

**Figure 2.**
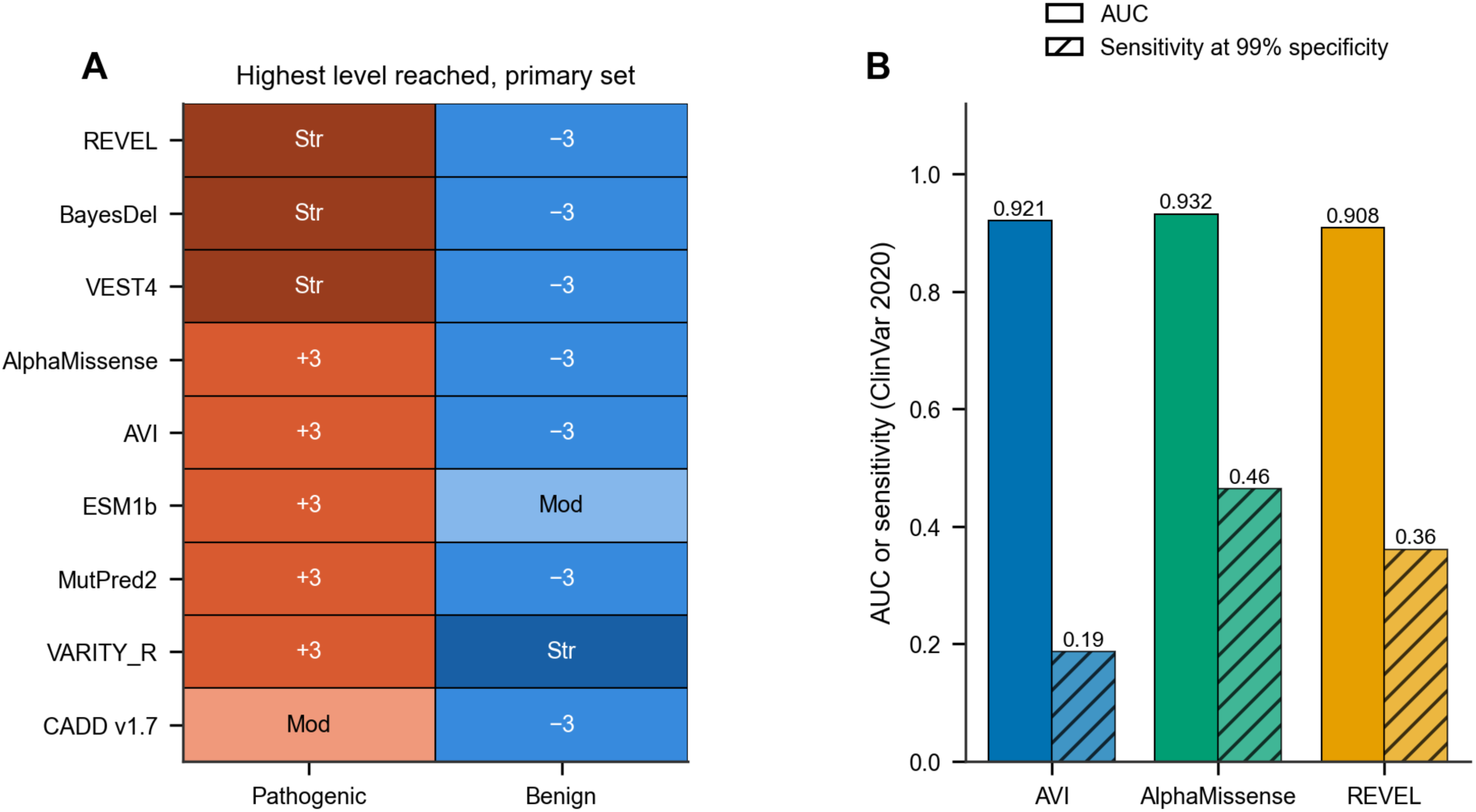
AVI calibration for missense variants. (A) Highest evidence level reached by AVI (AlphaGenome Variant Impact score) and eight other prediction tools in the primary set (rare variants without possible overlap with AVI training data; pathogenic levels in orange-red and benign levels in blue, darker for stronger evidence; Mod, moderate; Str, strong; +3 and −3, levels between moderate and strong; CADD v1.6 is shown in Table S2a). Levels whose threshold was reached by at most one labeled variant of their direction are not counted (Table S2a). AVI thresholds are given in Table 1. (B) Area under the receiver operating characteristic curve (AUC) and sensitivity at 99% specificity for AVI, AlphaMissense, and REVEL on ClinVar 2020 variants. Differences with 95% intervals are given in Table S7.

**Table 1.**
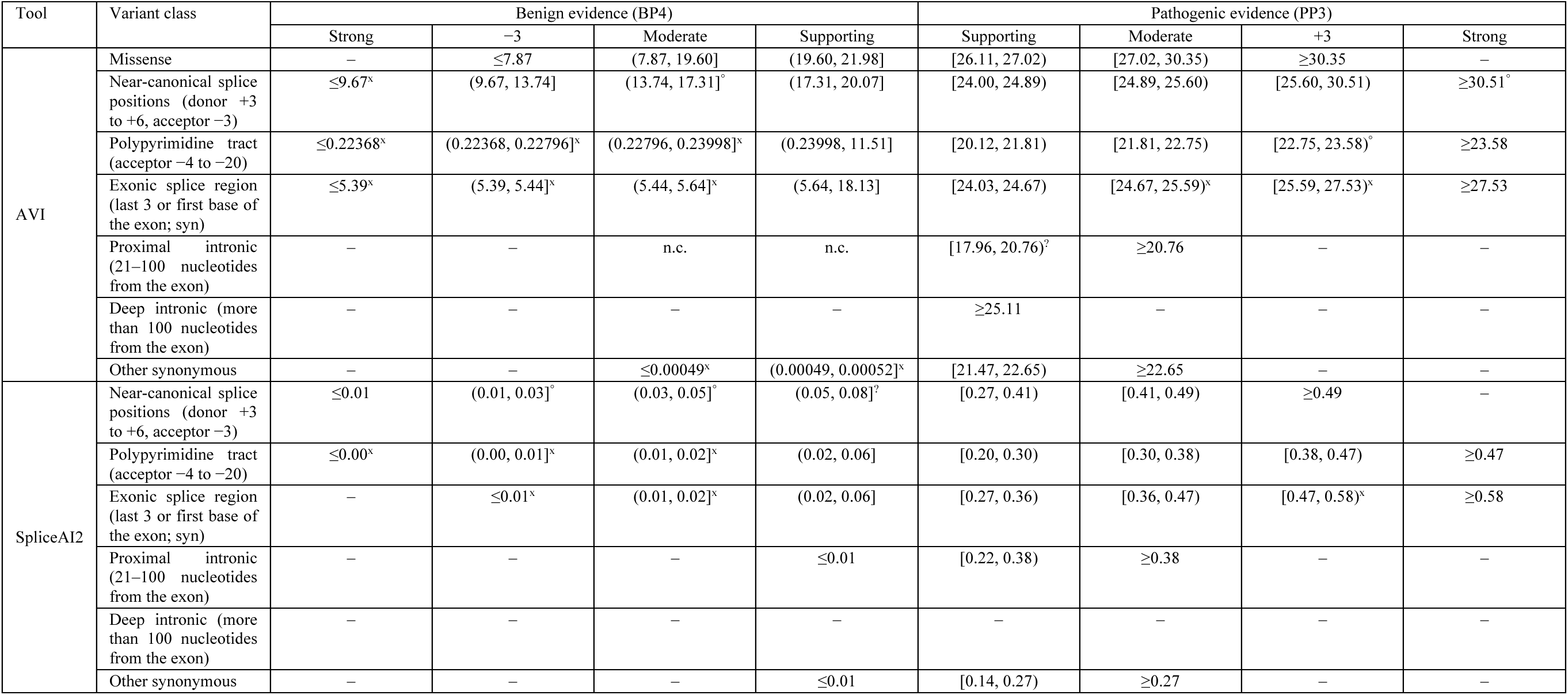
Score ranges of each evidence level in the main analysis. Each range runs from the threshold of its level to that of the next stronger level; thresholds are one-sided 95% bootstrap bounds, with ClinVar benign classifications as the benign class. Benign levels are on the left and pathogenic levels on the right, so scores increase from left to right. AVI scores are on the PHRED scale of the AVI release, rounded to two decimals toward the stricter side (values below 1 unrounded); SpliceAI2 scores are given in steps of 0.01. Benign ranges at the bottom of the score scale (AVI below 1, SpliceAI2 0.00–0.02) are separated by small score differences, so scores must be compared unrounded. Validation on later ClinVar classifications (ClinVar 2020 for missense variants, variants newly classified by September 2026 for the other classes; Table S3): no mark, validated; °, probably validated; ?, unclear; x, not tested (definitions in Material and methods). –, not reached, or not estimated for levels above moderate in proximal intronic and other synonymous variants (too few labeled variants); n.c., not counted (reached by at most one labeled variant); +3 and −3, evidence levels between moderate and strong; syn, synonymous; AVI, AlphaGenome Variant Impact score.

When rare population variants were used as controls instead of ClinVar benign variants, AVI reached pathogenic supporting and moderate evidence at stricter thresholds (Table S4a). The pathogenic +3 level could not be assessed in the uncorrected analysis because of the likelihood-ratio cap, and it was not reached after correction for an assumed pathogenic fraction of 4.41% (Table S4b). The benign −3 level was not reached.

Among the nine tools shown in Figure 2A, REVEL, BayesDel, and VEST4 reached strong pathogenic evidence; AVI, AlphaMissense, ESM1b, MutPred2, and VARITY_R reached +3 evidence; and CADD v1.7 reached moderate evidence (Table S2a; supplemental results). On ClinVar 2020 variants, AVI had a lower AUC than AlphaMissense (0.921 vs. 0.932; difference −0.012, 95% interval −0.017 to −0.006), whereas their AUCs were not distinguishable on ClinVar 2019 variants (0.932 vs. 0.927; difference 0.005, 95% interval −0.002 to 0.011). AVI also had lower sensitivity at 99% specificity than AlphaMissense on both releases (0.19 vs. 0.46 on ClinVar 2020, Figure 2B; 0.27 vs. 0.36 on ClinVar 2019). In the population reference set, AVI had high rank correlations with AlphaMissense (0.81) and CADD v1.7 (0.86), although its correlation with AlphaMissense was lower among pathogenic variants (0.49 in ClinVar 2019 and 0.50 in ClinVar 2020; Table S5). Adding AVI to AlphaMissense, which is already a component of AVI, did not improve the AUC of a two-score model. In contrast, adding AVI to REVEL increased the AUC by 0.029 (Tables S6a and S7).

### Both tools reached moderate evidence in five of six classes

Six non-coding and synonymous variant classes were included in the main analysis, with pathogenic variant counts ranging from 135 in proximal introns to 1,568 at near-canonical splice positions (Table S8). The donor flank and untranslated region classes contained too few pathogenic variants for calibration. The promoter class, included as an exploratory class, met the minimum sample-size requirements, but its AVI thresholds are reported separately in Table S9 and are not discussed further. For proximal intronic and other synonymous variants, sample sizes permitted calibration only up to moderate evidence (supplemental results).

AVI reached strong pathogenic evidence at near-canonical positions, in the polypyrimidine tract, and in the exonic splice region. It reached moderate evidence in proximal introns and at other synonymous sites, and supporting evidence in deep introns (Table 1; Figure 3; complete thresholds in Tables S9 and S10). SpliceAI2 reached the same levels, except at near-canonical positions, where it reached +3 rather than strong evidence, and in deep introns, where it reached no evidence level. Both tools nevertheless distinguished pathogenic from benign deep intronic variants (AUC 0.85 for each; Table S7). For SpliceAI2 in deep introns, the point estimate met the supporting requirement at 0.66, but 23% of bootstrap replicates did not, so the level was not reached. With a smaller window (1% instead of 3% of the reference set), supporting evidence was reached at 0.41 (Table S1). The missing level therefore reflects, at least in part, the window requirement rather than an absence of signal.

**Figure 3.**
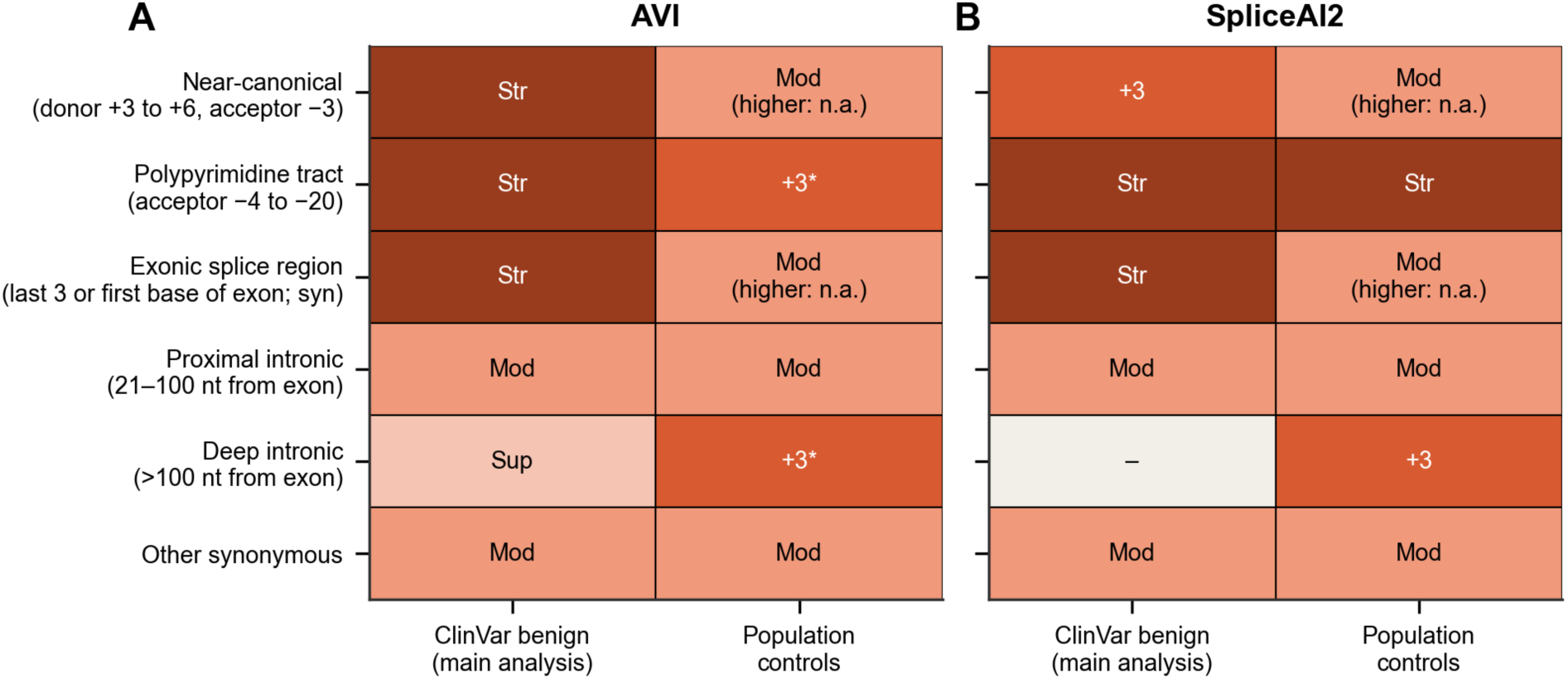
Highest pathogenic evidence levels across non-coding and synonymous classes. Highest levels reached by AVI (A) and SpliceAI2 (B) with ClinVar benign classifications (main analysis) and with rare population variants as controls. Darker colors indicate stronger evidence. AVI, AlphaGenome Variant Impact score; Sup, supporting; Mod, moderate; +3, level between moderate and strong; Str, strong; nt, nucleotides; syn, synonymous; –, no level reached; higher: n.a., levels above the one shown could not be assessed because of the likelihood-ratio cap in the uncorrected population-control analysis. Levels above moderate were not estimated for proximal intronic and other synonymous variants. *With population controls, the AVI strong-evidence threshold (34.76 in the polypyrimidine tract and 48.15 in deep introns) was at or above the highest pathogenic score in the class, so strong evidence was not counted. The highest evidence level reached should not be interpreted as a ranking of prediction tools.

Benign supporting evidence was reached in five classes by SpliceAI2 and four by AVI (Table 1). In proximal introns, the AVI benign thresholds were below all labeled variant scores and were therefore not counted as reached. At other synonymous sites, the AVI benign thresholds (0.00049 and 0.00052) were at the extreme lower end of the score distribution. Higher benign evidence levels are shown in Table 1 and detailed in Table S3.

### No threshold failed on later classifications

No evidence level reached by either tool failed validation on later ClinVar classifications, whether all newly classified variants or only first-time submissions were considered. Among the 35 pathogenic levels in Table 1, 29 were validated, two were probably validated, one was unclear, and three were not tested. Benign evidence levels were less extensively validated: among 30 levels, 12 were validated, three were probably validated, one was unclear, and 14 were not tested. When validation was restricted to first-time submissions, 21 of the 27 benign levels in non-coding and synonymous classes could not be tested (Table S3). The two unclear levels were AVI pathogenic supporting evidence in proximal introns and SpliceAI2 benign supporting evidence at near-canonical positions. Under the simpler rule of Pejaver et al., which requires only the likelihood-ratio point estimate to meet the required value, neither level would pass (2.33 vs. ≥2.406 and 0.417 vs. ≤0.416, respectively).^3^

### Moderate evidence was robust outside deep introns

In all five classes other than deep introns, both tools consistently reached pathogenic supporting and moderate evidence across robustness analyses in which calibration was possible, although the corresponding thresholds varied (Table S1; random-removal analyses in Tables S9 and S10). Even when class definitions remained unchanged, thresholds shifted by up to about one evidence level (supplemental results). Resampling variants instead of genes, as in the missense analysis, gave the same levels in five of the six classes; in deep introns, it also reached benign supporting evidence for both tools and pathogenic supporting evidence for SpliceAI2 (at 0.68; Table S1), consistent with the clustering of labeled deep intronic variants within genes.

Deep intronic thresholds were substantially less stable. Across the 13 robustness analyses in Tables S9 and S10 that kept the class definitions unchanged, the AVI supporting threshold ranged from 17.13 to 25.57, and AVI also reached moderate evidence in four analyses (thresholds 24.10–33.75). SpliceAI2 reached supporting evidence in six of the 13 analyses, with thresholds ranging from 0.26 to 0.68.

The 100-nucleotide boundary between proximal and deep introns is a convention, so we also tested boundaries at 200 and 500 nucleotides. With these boundaries, both tools still reached supporting and moderate evidence in proximal introns, but the moderate thresholds rose to as high as 23.82 for AVI and 0.70 for SpliceAI2. In deep introns, only AVI reached a pathogenic evidence level: supporting evidence at 37.48 with the 200-nucleotide boundary, a score that no variant with a later classification reached. With the 500-nucleotide boundary, AVI reached only benign evidence levels (Table S1).

To assess whether differences in allele frequency between pathogenic and benign variants affected the thresholds, we restricted both ClinVar classes to variants absent from gnomAD v4.1 genomes; this changed pathogenic supporting and moderate thresholds by no more than 0.57 for AVI and 0.02 for SpliceAI2 outside proximal and deep introns. In proximal introns, thresholds increased by 1.14–1.51 for AVI and by 0.06–0.07 for SpliceAI2. In deep introns, the AVI supporting threshold increased from 25.11 to 33.21, although the point estimate on which it is based changed only slightly (22.09 to 22.81; Tables S9 and S11). At near-canonical positions, this restriction halved the benign class (1,406 to 703 variants) and increased the AVI +3 threshold from 25.60 to 27.62, despite little change in the point estimate (25.14 to 24.91). AVI strong evidence was no longer reached (Table S11).

### Population controls changed pathogenic evidence thresholds

With rare population variants as controls, both tools reached supporting and moderate pathogenic evidence in all six classes (Table S4a). However, the thresholds differed from those of the main analysis. In proximal and deep introns, population controls produced lower thresholds, and both tools reached +3 evidence in deep introns (Figure 3), compared with supporting evidence for AVI and no reached level for SpliceAI2 in the main analysis. In the three classes adjacent to exons and at other synonymous sites, population controls produced higher thresholds.

Among these four classes, evidence above moderate could be assessed in the uncorrected analysis only in the polypyrimidine tract (at near-canonical positions and in the exonic splice region, these levels were not assessable, and at other synonymous sites, they were not estimated). There, both tools reached +3 evidence at higher thresholds, and SpliceAI2 also reached strong evidence. The SpliceAI2 strong level stands, because the likelihood-ratio cap can prevent a level from being reached but cannot produce it. For AVI, the strong-evidence threshold (34.76) lay at or above the highest pathogenic score in the class, so strong evidence was not counted.

As AVI was trained on allele frequency, we also used higher-frequency population variants as controls; these gave thresholds similar to those obtained with rare population variants in proximal and deep introns (Table S12a). Near exons, few higher-frequency variants were available (276 at near-canonical positions and 1,630 in the polypyrimidine tract); the 45 in the exonic splice region were too few for calibration. As thresholds from small sets are less precise, we compared the near-exon thresholds with thresholds from 20 random sets of the same size drawn from ClinVar benign classifications and from rare population variants. In the polypyrimidine tract, thresholds obtained with higher-frequency controls fell within the ClinVar-based range for both tools. At near-canonical positions, AVI thresholds fell within or at the upper edge of the ClinVar-based range and below all thresholds from rare population variants, whereas SpliceAI2 thresholds were above the ClinVar-based range and closer to the rare-population range (Table S12b).

### Likely benign classifications changed abruptly at 20 nucleotides

We compared ClinVar classifications with splicing-assay results at each intronic position. The number of likely benign classifications differed by 64-fold between acceptor positions −20 and −21 and by 62-fold between donor positions +20 and +21. In comparison, benign classifications differed by only 4-and 5-fold, respectively. The proportion of splice-disruptive substitutions in the splicing assay remained similar across these boundaries (10.0% vs. 10.4% and 1.9% vs. 2.0%; Figure 4A; Table S13).

**Figure 4.**
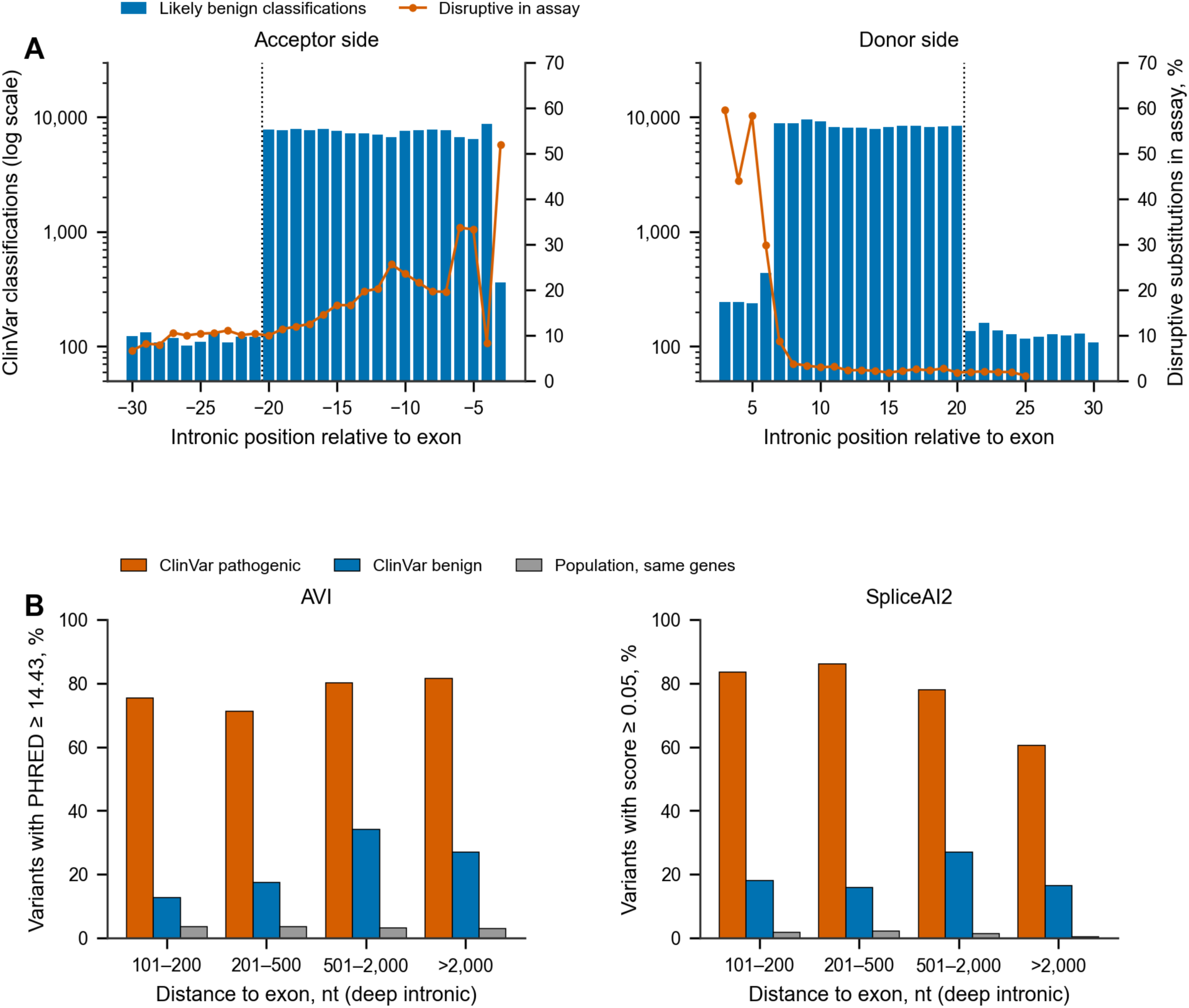
Benign classifications by intronic position and scores of deep intronic variants. (A) Number of likely benign ClinVar classifications and proportion of splice-disruptive substitutions in the splicing assay by intronic position. Dotted lines mark the boundary between positions 20 and 21 nucleotides from the exon. (B) Proportion of ClinVar pathogenic, ClinVar benign, and population variants from the same genes scoring at or above the pathogenic supporting threshold obtained with population controls, stratified by distance from the exon in deep introns. AVI scores are on the PHRED scale; nt, nucleotides; AVI, AlphaGenome Variant Impact score.

### Benign deep intronic variants scored higher than population variants

To understand why ClinVar benign classifications and population controls gave different deep intronic thresholds, we compared their scores. In deep introns, ClinVar benign variants had substantially higher prediction scores than rare population variants. At the AVI supporting threshold obtained with population controls (14.43), 25% of ClinVar benign variants scored at or above the threshold, compared with 3.0% of population variants. At the corresponding SpliceAI2 threshold (0.05), the proportions were 20% and 1.0%. These differences persisted after stratification by distance from the exon and restricting population variants to the same genes, ranging from 4-to 11-fold for AVI and from 7-to 35-fold for SpliceAI2 (Figure 4B; Table S14). By contrast, differences between ClinVar benign and population variants were smaller in proximal introns (1.4-to 4.7-fold; Table S14).

These high-scoring benign variants were no more concentrated in a few genes than deep intronic benign variants overall (Table S15; supplemental results).

### Thresholds held on classifications made before SpliceAI

Classifications made after SpliceAI was published may have used its predictions, so we separated those made before it. Among calibration variants, 22% of pathogenic and 6% of benign variants (734 and 32,357, respectively) already carried their classifications in December 2018. When calibration was repeated using only the later classifications, pathogenic thresholds reached in both calibrations differed by up to 1.03 for AVI and 0.03 for SpliceAI2 outside deep introns. In deep introns, the AVI supporting threshold increased from 25.11 to 32.08 (Table S16a).

When these later-calibrated thresholds were evaluated against the earlier classifications, none of the 29 pathogenic evidence levels failed: 11 were validated, eight were probably validated, one was unclear (AVI moderate evidence in the polypyrimidine tract), and nine were not tested. Among 22 benign evidence levels, none failed: nine were validated, one was probably validated, one was unclear, and 11 were not tested (Table S16b).

In every class with a main threshold, scores at or above the main supporting threshold gave likelihood ratios well above the supporting requirement, on both earlier and later classifications. For AVI in deep introns, these likelihood ratios were 22 and 15, respectively (Table S16c). Furthermore, outside deep introns, the AUC difference between SpliceAI2 and AVI did not change detectably between earlier and later classifications.

In deep introns, however, the difference in AUC shifted toward SpliceAI2 by 0.14 (95% interval 0.05–0.25). On earlier classifications, the AUCs were 0.89 for AVI and 0.78 for SpliceAI2 (76 pathogenic and 304 benign variants); on later classifications (434 pathogenic and 3,586 benign variants), they were 0.84 and 0.86 (Table S16d).

### The same score provided different evidence across variant classes

To test whether one calibration could serve all intronic classes, we also calibrated the four intronic classes together. The resulting AVI score intervals did not provide equivalent evidence across classes. At near-canonical positions, the supporting and moderate intervals had likelihood ratios of 0.81 (95% interval 0.57–1.15) and 2.5, respectively, below the required values of 2.406 and 5.790 (Table S17). In deep introns, the likelihood-ratio point estimate for the pooled moderate interval was also below its requirement.

For SpliceAI2, by contrast, the likelihood-ratio point estimates of the pooled pathogenic intervals met their requirements in all four classes, although the likelihood ratio of the same interval still differed between classes, for example from 5.8 at near-canonical positions to 54 in proximal introns for the moderate interval. However, its pooled benign supporting interval gave too little benign evidence in proximal and deep introns (likelihood ratios 0.63 and 0.57; required maximum 0.416). Its pooled benign moderate interval also gave too little benign evidence in deep introns (0.28 vs. ≤0.173; Table S17).

### Prior probabilities differed across classes

The likelihood-ratio requirements assume a prior probability of 4.41%, so we estimated the prior range in each class. The proportion of rare population variants already classified as pathogenic ranged from 0.90% in the exonic splice region to 0.00033% in deep introns. The estimated upper limit for the prior probability of pathogenicity ranged from approximately 13% to below 0.1% (Table 2). The 4.41% prior fell within the estimated range only at near-canonical positions and in the exonic splice region; it exceeded the upper limit in the other four classes.

**Table 2.** Estimated range of prior pathogenicity probabilities among rare population variants and corresponding posterior probabilities for each evidence level. Lower and upper estimates define a range for the prior probability of pathogenicity. The upper limit (ceiling) is the smallest of nine upper confidence limits (one at each decile of the pathogenic scores), computed separately from the scores of each tool; it assumes that undiscovered pathogenic variants have score distributions similar to those of known pathogenic variants. Posterior ranges use the floor and the higher of the two upper limits. Mixture-proportion estimates were obtained using five prediction scores other than AVI and SpliceAI2 and exceeded the estimated upper limits in proximal introns, deep introns, and at other synonymous sites. The reported posterior probabilities are not intended for direct use in points-based evidence calculations. Probabilities are calculated at the minimum likelihood ratio of each level. AVI, AlphaGenome Variant Impact score; LR, likelihood ratio; floor, share of rare population variants already classified as pathogenic or likely pathogenic; syn, synonymous.

| Variant class | Known pathogenic (floor), % | Ceiling, upper limit (AVI / SpliceAI2), % | Mixture estimate, % (95% interval) | Probability of pathogenicity at supporting (LR 2.406), % | at moderate (LR 5.790), % | at strong (LR 33.53), % |
| --- | --- | --- | --- | --- | --- | --- |
| Near-canonical splice positions (donor +3 to +6, acceptor -3) | 0.50 | 8.5 / 11 | 5.4 (4.2–6.7) | 1.2–23 | 2.8–42 | 14–81 |
| Polypyrimidine tract (acceptor -4 to -20) | 0.069 | 1.7 / 2.1 | 1.6 (1.4–1.9) | 0.16–5.0 | 0.40–11 | 2.2–42 |
| Exonic splice region (last 3 or first base of the exon; syn) | 0.90 | 13 / 12 | 8.9 (6.1–12) | 2.1–26 | 5.0–46 | 23–83 |
| Proximal intronic (21–100 nucleotides from the exon) | 0.0026 | 0.27 / 0.19 | 1.4 (1.2–1.7) | 0.0062–0.65 | 0.015–1.6 | 0.086–8.4 |
| Deep intronic (more than 100 nucleotides from the exon) | 0.00033 | 0.083 / 0.055 | 1.6 (1.4–2.0) | 0.00079–0.20 | 0.0019–0.48 | 0.011–2.7 |
| Other synonymous | 0.0074 | 0.34 / 0.33 | 1.7 (1.3–2.2) | 0.018–0.82 | 0.043–1.9 | 0.25–10 |

If these upper limits are valid, supporting and moderate evidence would correspond to probabilities of pathogenicity about 20 times lower or more in proximal and deep introns and at other synonymous sites than at near-canonical positions and in the exonic splice region. Mixture-proportion estimates, a third estimate of the pathogenic fraction, were consistent with the estimated ranges in the three classes adjacent to exons but exceeded them in the other three classes. This discrepancy suggests either that the mixture method cannot reliably estimate pathogenic fractions far below 1%, or that undiscovered pathogenic variants tend to receive lower scores than known pathogenic variants. In that case, the upper limits would be too low, but the mixture estimates would be unreliable as well, because the mixture method also assumes that known pathogenic variants represent undiscovered ones.

When the likelihood-ratio requirements were recalculated for priors down to 0.5%, the supporting and moderate thresholds in the polypyrimidine tract and at other synonymous sites rose by at most 1.45 for AVI and 0.12 for SpliceAI2 (Tables S9 and S10). The moderate thresholds in proximal introns rose to 24.12 for AVI and 0.81 for SpliceAI2, and AVI no longer reached supporting evidence in deep introns at priors of 1% or lower. Even a prior of 0.5% exceeds the estimated upper limits in proximal and deep introns and at other synonymous sites.

### SpliceAI2 had a higher AUC in two of six classes

Using ClinVar classifications, the AUCs of AVI and SpliceAI2 did not differ detectably in four of the six classes. SpliceAI2 had higher AUCs in the polypyrimidine tract (0.991 vs. 0.977) and proximal introns (0.927 vs. 0.896) when ClinVar benign variants were used as the negative class. This advantage was also observed when population variants were used as controls (Figure 5; Table S7). In three of the six classes, the AUCs of the original SpliceAI (±4,999-nucleotide window) and SpliceTransformer differed from those of AVI and SpliceAI2 by less than 0.03 (Table S18).

**Figure 5.**
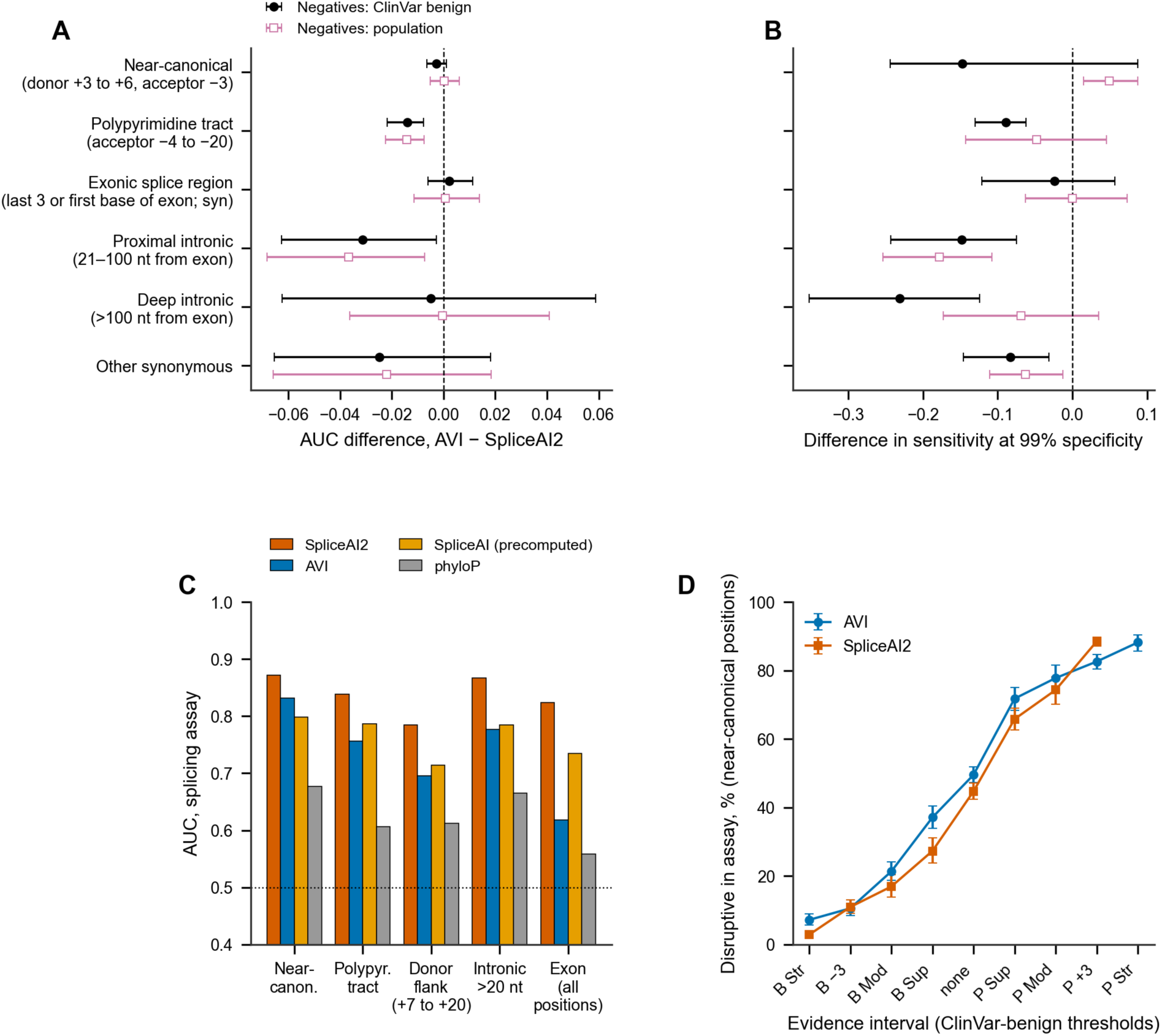
Comparison of AVI and SpliceAI2. (A) Differences in the area under the receiver operating characteristic curve (AUC; AVI minus SpliceAI2) with 95% gene-cluster bootstrap intervals, by variant class and negative class. (B) Differences in sensitivity at 99% specificity (AVI minus SpliceAI2) with 95% gene-cluster bootstrap intervals. In A and B, values below 0 favor SpliceAI2. (C) AUCs in the splicing assay by position class, on substitutions scored by all four tools (AVI, SpliceAI2, the original SpliceAI as precomputed scores, and phyloP; Table S20); Near-canon., near-canonical; Polypyr., polypyrimidine; >20 nt, intronic positions more than 20 nucleotides from the exon; dotted line, AUC 0.5; the vertical axis starts at 0.4. (D) Proportion of splice-disruptive substitutions at near-canonical positions across calibrated evidence intervals, with 95% intervals; B and P, benign and pathogenic intervals; none, no evidence; Sup, supporting; Mod, moderate; +3 and −3, levels between moderate and strong; Str, strong; nt, nucleotides; syn, synonymous; AVI, AlphaGenome Variant Impact score.

At 99% specificity, SpliceAI2 had higher sensitivity in four classes with ClinVar benign variants and in two with population variants as the negative class. By contrast, AVI had higher sensitivity at near-canonical positions when population variants were used as controls. In two-score models, adding SpliceAI2 to AVI improved AUC in five of the six classes. Adding AVI to SpliceAI2 increased the AUC slightly at near-canonical positions and in the exonic splice region, and increased AUC by 0.029 in deep introns when population variants were used as controls (Table S6a).

To explore whether AVI adds information beyond splicing prediction, for example through predicted effects on promoters, enhancers, polyadenylation, or RNA abundance, we counted pathogenic variants that reached an AVI pathogenic threshold without a predicted splicing effect (SpliceAI2 score below 0.05). Only 18 of 3,277 (0.5%) did so, ten of them in deep introns. Outside deep introns, none of the 658 pathogenic classifications made before SpliceAI did so (Table S6b).

To relate our results to current practice, we also evaluated the original SpliceAI at the cutoff proposed by Walker et al. With a ±4,999-nucleotide window, scores ≥0.2, taken together, had likelihood ratios of 7 to 36 across classes (Table S19a). These are averages over the whole category, which includes much higher scores. In contrast, class-specific calibration of the original SpliceAI placed its supporting-evidence threshold between 0.24 and 0.45 in five classes and at 0.79 in deep introns (Table S19b).

### SpliceAI2 outperformed AVI in the splicing assay

To compare the tools without relying on clinical classifications, we used a splicing assay that measured exon inclusion for 310,405 substitutions. For intronic substitutions outside the canonical splice-site dinucleotides, SpliceAI2 achieved an AUC of 0.90 in the splicing assay, compared with 0.84 for AVI, 0.84 for the original SpliceAI (precomputed scores, 50-nucleotide window), and 0.71 for phyloP (Table S20). SpliceAI2 had higher AUC than AVI in every position class outside the canonical dinucleotides, with differences ranging from 0.04 at near-canonical positions to 0.21 in exons (Figure 5C; Table S7). The exonic class included missense substitutions, whose AVI scores also reflect predicted effects on the protein, so this comparison favors a dedicated splicing model.

To check whether stronger evidence corresponds to more splice disruption, we placed each near-canonical substitution in the assay into the evidence interval that contains its score. The proportion of splice-disruptive substitutions increased from 7% in the strongest benign interval to 88% in the strongest pathogenic interval for AVI, and from 3% to 89% for SpliceAI2. Within the benign supporting interval, 37% (AVI) and 27% (SpliceAI2) of substitutions were disruptive, compared with 49% of all substitutions at these positions (Figure 5D; Table S21; supplemental results).

## Discussion

### What the two scores support

For missense variants, AVI reached pathogenic evidence up to +3 and benign evidence up to −3. For non-coding and synonymous variants, both AVI and SpliceAI2 reached pathogenic supporting and moderate evidence in every class except deep introns, where AVI reached supporting evidence and SpliceAI2 no level. Of the 23 pathogenic supporting and moderate levels in Table 1, 21 were validated on later ClinVar classifications, one was unclear, one was not tested, and none failed. Moreover, these levels were reached with rare population variants as controls, although stricter thresholds were required for missense variants, in the three classes adjacent to exons, and at other synonymous sites.

By contrast, higher evidence levels were less robust. With population controls in the uncorrected analysis, they could not be assessed at near-canonical positions or in the exonic splice region, and AVI strong evidence in the polypyrimidine tract was not counted. For missense variants, AVI pathogenic +3 evidence could not be assessed with population controls and was not reached after correction for an assumed pathogenic fraction of 4.41%.

Thus, supporting and moderate evidence could be reached with either comparison group, but the thresholds differed substantially. In the exonic splice region, for example, the supporting threshold obtained with population controls was higher than the +3 threshold of the main analysis for AVI and higher than the strong threshold for SpliceAI2. The main thresholds are given in Table 1 and those obtained with population controls in Tables S4a and S4b, to show how much they depend on this choice.

### Benign classifications limit non-coding calibration

In ClinVar, the number of likely benign classifications changed approximately 60-fold at the boundary 20 nucleotides from the exon, whereas the proportion of splice-disruptive substitutions in the splicing assay remained similar on either side. The preferential reporting of intronic variants close to exons is well recognized;^9,33^ the splicing assay adds direct evidence that this boundary reflects reporting practices rather than a change in splicing.

Farther from exons, ClinVar benign variants also differed from typical population variants. In deep introns, benign-classified variants exceeded the pathogenic thresholds from population-control calibration much more often than population variants from the same genes and at comparable distances from the exon. This difference was not explained by a few highly represented genes (Table S15).

Two technical explanations remain possible. Variant classes were defined on MANE Select transcripts, so some deep intronic variants may lie in exons or splice regions of other transcripts. In addition, population variants had to pass gnomAD quality filters, whereas ClinVar variants did not, so the two groups may differ in which genomic positions they include, and therefore in their scores. Beyond these technical explanations, the gene set included all genes with any pathogenic classification, so some benign variants may alter splicing where this is tolerated, for example in genes without a loss-of-function mechanism.

Near exons, by contrast, population controls gave higher thresholds than ClinVar benign classifications. Undiscovered pathogenic variants among the controls explain only part of this difference. With a correction assuming that 10% of the controls were pathogenic, near or above the estimated upper limits in these classes (Table 2), the supporting and moderate thresholds of both tools remained higher than those obtained with ClinVar benign classifications in the three classes adjacent to exons (Table S1). Higher-frequency population variants gave AVI thresholds within or at the upper edge of the ClinVar-based range at equal sample sizes, but this analysis is not independent of AVI. SpliceAI2 thresholds from the same variants were above this range at near-canonical positions and within it in the polypyrimidine tract. Benign classifications near exons and at synonymous sites may themselves rest on splicing predictions (BP4, or BP7 for synonymous variants and for intronic variants at +7/−21 or beyond),^8^ which would make the ClinVar-based thresholds too lenient.

Neither ClinVar benign classifications nor rare population controls can be assumed to represent the true benign distribution. Undiscovered pathogenic variants bias likelihood ratios from population controls toward 1, whereas ClinVar benign variants are selected during clinical testing and reporting. Because ClinVar benign classifications, rare population variants, and higher-frequency population variants are selected in different ways, agreement between any two of them does not establish which thresholds are correct. Our analyses cannot determine the cause of these differences.

### AVI and SpliceAI2 compared with existing tools

Overall, AVI did not outperform AlphaMissense for missense variants or SpliceAI2 in the non-coding and synonymous classes. For missense variants, its AUC was lower than that of AlphaMissense in ClinVar 2020 but did not differ detectably in ClinVar 2019, while its sensitivity at 99% specificity was lower in both releases. Adding AVI to AlphaMissense did not improve predictive performance, which is unsurprising given that AlphaMissense is already one of AVI’s input features. This suggests that AVI’s additional features provide limited incremental information for missense variant classification. Adding AVI to REVEL increased the AUC by 0.029, but this gain may come from its AlphaMissense component, which alone had a higher AUC than REVEL on ClinVar 2020 (0.932 vs. 0.908; Tables S6a and S7).

In the six non-coding and synonymous classes, the AUCs of AVI and SpliceAI2 did not differ detectably in four. SpliceAI2 had higher AUC in two classes and higher sensitivity at 99% specificity in four classes with ClinVar benign variants as the negative class, although AVI had the higher AUC on deep intronic classifications made before SpliceAI. SpliceAI2 also outperformed AVI in every position class of the splicing assay outside the canonical dinucleotides. The assay does not depend on clinical classifications, but the SpliceAI2 developers used it to evaluate their model.^13^

However, the comparison between AVI and SpliceAI2 was not fully matched for potential training-data overlap: variants potentially associated with AVI training were excluded, whereas variants in genes used to train SpliceAI2 were not, which, if anything, favors SpliceAI2 (see Limitations). Moreover, the highest evidence level reached (Figures 2A and 3) is not a direct measure of predictive performance, because it also depends on score resolution and the window requirements of the calibration method. In addition, the two tools serve different purposes. SpliceAI2 predicts only splicing effects, whereas AVI scores coding and non-coding variants on one scale from many predicted molecular effects and shows which effect drives each score,^12^ which few tools offer.

### The importance of class-specific thresholds

The same prediction score did not provide equivalent evidence across variant classes: when the four intronic classes were calibrated together, the AVI supporting and moderate intervals provided less than the required evidence at near-canonical positions, and the moderate point estimate fell short in deep introns. A recent preprint found a similar pattern for AVI levels derived for intronic and untranslated variants together.^14^

For SpliceAI2, by contrast, the point estimates of the pooled pathogenic intervals met their requirements in all intronic classes, but the evidence within the same interval still varied several-fold between classes (Table S17), and the pooled benign intervals provided too little benign evidence in proximal and deep introns. These findings support class-specific calibration; the pooled thresholds were descriptive and are not proposed for use.

### Evidence levels and previous work

The evidence levels in this study are defined by likelihood ratios. Likelihood ratios do not depend on the prior probability of pathogenicity,^3^ but the resulting posterior probabilities do. As in earlier calibrations that used a common prior for all missense variants^3^ or splicing positions,^8^ we kept the same level names across classes; Table 2 shows how the posterior probability of each level changes across the estimated class-specific priors.

However, the commonly used 4.41% prior fell within our estimated range only at near-canonical positions and in the exonic splice region; in the other four classes, the estimated upper limit was about 2-fold to more than 50-fold lower. Consequently, the same evidence level may correspond to a much lower probability of pathogenicity in these classes, and applying it unchanged in a points-based framework whose evidence strengths were set for a 10% prior^2^ and, for computational evidence, a 4.41% prior^3^ could overestimate that probability. These upper limits, however, assume that undiscovered pathogenic variants score like known ones, and they describe rare population variants, whereas variants found in diagnostic testing are generally enriched for pathogenic variants.

Finally, our results are compatible with the SpliceAI cutoff of 0.2 recommended by the ClinGen SVI Splicing Subgroup,^8^ which was derived for altered splicing rather than pathogenicity and with different evidence-strength requirements. As a single category, SpliceAI scores ≥0.2 provided at least moderate evidence in every class, but in our class-specific calibration of SpliceAI, the supporting thresholds were 0.24 to 0.45 in five classes and 0.79 in deep introns (Table S19b), so scores just above 0.2 did not meet the supporting requirement on their own. Evidence for the whole category should therefore not be assumed to apply to every score within it.

### Deep intronic variants

Deep intronic variants were the hardest to calibrate. In the main analysis, only AVI reached a pathogenic evidence level (supporting, at 25.11), although both tools distinguished pathogenic from benign deep intronic variants (AUC 0.85) and both reached +3 evidence with population variants as controls.

Although the main AVI threshold was validated on later ClinVar classifications, it was highly sensitive to analytical choices. The supporting threshold ranged from 17.13 to 25.57 across 13 robustness analyses with unchanged class definitions (Tables S1 and S9) and increased to 33.21 when both ClinVar classes were restricted to variants absent from gnomAD (Table S11). Moreover, moving the proximal–deep intronic boundary to 200 nucleotides increased the threshold to 37.48, which no variant in the later validation set reached. With a 500-nucleotide boundary, AVI no longer reached pathogenic evidence (Tables S1 and S9). Similarly, SpliceAI2 reached supporting evidence in only six of the 13 robustness analyses (Tables S1 and S10).

These findings also show a limitation of the calibration method. Each score window must contain at least 3% of the reference set, but in deep introns only about 1%–3% of reference variants scored at or above the pathogenic supporting threshold obtained with population controls. The method may therefore be unreliable for classes in which most variants have low scores.

The clinical labels introduce further uncertainty. ClinVar benign deep intronic variants may not be representative of population variation, and computational predictions may have contributed to the identification of known pathogenic variants. The AUC difference between SpliceAI2 and AVI shifted by 0.14 in favor of SpliceAI2 between earlier and later classifications. However, the earlier set included only 76 pathogenic variants, limiting the precision of this comparison. Population controls replace only the benign class and cannot correct selection among pathogenic variants.

Together, these findings indicate that the deep intronic AVI threshold is provisional. Calibration methods adapted to score distributions concentrated near zero, and less selectively ascertained variant classifications, are needed before deep intronic evidence can be assigned with confidence.

### Limitations

ClinVar classifications of non-coding and synonymous variants may have incorporated computational splicing predictions. Nevertheless, outside deep introns we found no clear sign that this influenced the thresholds: thresholds derived from later classifications were similar to the main thresholds and did not fail on classifications made before SpliceAI. Furthermore, later classifications in these classes showed no detectable shift in relative performance toward SpliceAI2. However, benign classifications of synonymous and some intronic variants can rely on splicing predictions (BP7), which may inflate the apparent performance of splicing predictors in these classes.

Even so, these analyses cannot exclude prediction-related bias. The earlier sets were relatively small, leaving approximately one-third of pathogenic and half of benign evidence levels not tested, and they included only 2018 classifications that remained unchanged in 2025, excluding those that were later contested or revised. In addition, earlier classifications may have used splicing predictors that were available before 2019, and comparing AVI with SpliceAI2 would not detect biases that favor both tools equally.

Validation outcomes also need careful interpretation. Probably validated and unclear levels were not confirmed by later classifications; these verdicts show only that the level did not fail. In addition, many levels, particularly benign ones, could not be tested, and when later classifications include tens of thousands of benign but few pathogenic variants, benign levels can rarely fail validation. A pathogenic level could fail only if the 95th percentile of its likelihood ratio fell below the requirement, and a benign level only if the 5th percentile lay above it. Both are unlikely when few later pathogenic variants are available, so validation shows mainly that no threshold was clearly contradicted. We also did not adjust for multiple comparisons.

In addition, SpliceAI2 was trained on RNA sequencing data without clinical variant labels, holding out only genes on chromosomes 21 and 22,^13^ so most variants in our study lie in genes whose sequences were seen during training.

The splicing assay measures exon inclusion in a minigene system rather than pathogenicity and captures only effects on splicing, whereas AVI also incorporates predictions of chromatin accessibility, transcription-factor binding, histone modifications, transcription initiation, RNA abundance, polyadenylation, and chromatin interactions.^10,12^ The assay therefore cannot show whether AVI adds predictive value for variants acting through these other mechanisms, including promoter variants. Among known pathogenic variants, however, AVI rarely reached a pathogenic threshold without a predicted splicing effect, although variants acting through other mechanisms may be underrepresented in ClinVar. At near-canonical positions, about a third of assay substitutions in the benign supporting interval disrupted splicing, so benign evidence at these positions should be weighed against functional data when available. In addition, too few pathogenic variants were available to calibrate the donor flank and untranslated region classes.

Our incremental-value analyses used linear logistic models and may underestimate information that more complex models could capture. In addition, AVI was trained with allele frequency as its label, which could bias its scores. Restricting both ClinVar classes to variants absent from gnomAD genomes changed the AVI pathogenic supporting and moderate thresholds by at most 0.57 outside proximal and deep introns. In proximal introns, the thresholds of both tools rose, and in deep introns the AVI supporting threshold rose from 25.11 to 33.21. The point estimates changed less (in deep introns, from 22.09 to 22.81), so these increases mainly reflect the smaller sets; this analysis could therefore neither show nor exclude an effect of allele frequency on AVI.

Rarity was defined from gnomAD genome data, so in and near exons some variants absent from the genomes may still be common in the larger exome data. However, in a robustness analysis of missense variants, defining rarity jointly from exome and genome data changed AVI thresholds by no more than 0.05 (Table S2c).

Finally, few higher-frequency population variants were available near exons, and most were at least three times more frequent in one ancestry group than overall (Table S12c). Moreover, because variants already in ClinVar were excluded from the population reference set, these variants may not represent common population variation.

### Scope of the thresholds

The proposed thresholds are those of the main analysis (Table 1), because they follow the method of previous ClinGen calibrations. Thresholds obtained with population controls (Tables S4a and S4b), which were stricter near exons and at other synonymous sites, are given for comparison. Validation on later ClinVar classifications cannot show which of the two sets of thresholds is more accurate, because the later benign classifications may have been selected in the same way as the earlier ones. The thresholds apply to rare SNVs (absent from gnomAD v4.1 genomes or with a filtering allele frequency below 0.001); for the non-coding and synonymous classes, they apply to variants in genes with known pathogenic variants, with classes defined on MANE Select transcripts. Each variant receives the evidence level of the score range that contains its score; variants scoring between the benign and pathogenic supporting thresholds, or without a score, receive no computational evidence.

No thresholds are proposed for the donor flank (+7 to +20), canonical splice-site positions, untranslated regions, promoters (an exploratory class), or splicing effects of missense variants. The two tools were calibrated separately, so their evidence should not be combined for the same variant. In the polypyrimidine tract, in proximal and deep introns, and at other synonymous sites, the 4.41% prior exceeded the estimated upper limit, so the same level may correspond to a lower probability of pathogenicity than assumed (Table 2). In addition, the deep intronic AVI threshold was unstable across analyses and is provisional.

These thresholds are research calibrations, and AVI and SpliceAI2 are provided for research use only, according to the AlphaGenome Atlas^12^ and the SpliceAI2 license (Web resources). The thresholds also require replication in independent data and review by the relevant variant-classification expert groups before clinical implementation. Moreover, because they depended strongly on the choice between ClinVar benign classifications and population controls, future calibrations of non-coding evidence should report thresholds derived using both.

## Supporting information

Supplemental Methods and Results

## Data and code availability

Analysis code is available at Zenodo (10.5281/zenodo.23494356).

## Supplemental information

Supplemental information includes supplemental methods, supplemental results, and Tables S1–S22.

## Acknowledgments

We thank the developers of AlphaGenome and SpliceAI2 for making their precomputed scores available, the gnomAD and ClinVar teams and all ClinVar submitters for sharing their data, and Pejaver and colleagues for publishing their calibration code. Cloud computing was supported by free credits provided by Modal.

## Author contributions

Conceptualization, M.A.E., A.B., and Y.D.; methodology, M.A.E., A.B., and Y.D.; software, A.B.; formal analysis, A.B. and Y.D.; investigation, A.B., Y.D., C.F., T.D., and O.V.; validation, Y.D., C.F., T.D., and O.V.; data curation, A.B. and Y.D.; visualization, A.B.; writing – original draft, A.B. and Y.D.; writing – review & editing, all authors; supervision, M.A.E.

## Declaration of generative AI and AI-assisted technologies in the manuscript preparation process

During the preparation of this manuscript, the authors used Claude Opus 5.5 (Anthropic) to assist with improving the grammar and clarity of the text. The authors subsequently reviewed and edited all content to ensure accuracy and take full responsibility for the final manuscript.

## Web resources

AlphaGenome Atlas, https://alphagenome.google/atlas CADD, https://cadd.gs.washington.edu/

ClinGen calibration code (Pejaver et al.), https://github.com/pejaverlab/clingen-svi-comp_calibration

ClinVar, https://www.ncbi.nlm.nih.gov/clinvar/ Ensembl VEP REST API, https://rest.ensembl.org/ GENCODE, https://www.gencodegenes.org/ gnomAD, https://gnomad.broadinstitute.org/

MANE, https://www.ncbi.nlm.nih.gov/refseq/MANE/ OpenSplice, https://results.hgi.sanger.ac.uk/OpenSplice/

phyloP (UCSC Cactus 241-way), https://hgdownload.soe.ucsc.edu/goldenPath/hg38/cactus241way/

SpliceAI, https://github.com/Illumina/SpliceAI SpliceAI2, https://github.com/Illumina/SpliceAI2

SpliceAI2 precomputed scores, https://huggingface.co/datasets/illumina-ai/SpliceAI2-data

SpliceTransformer, https://github.com/ShenLab-Genomics/SpliceTransformer

Zenodo, https://zenodo.org/

## References

1. Richards, S., Aziz, N., Bale, S., Bick, D., Das, S., Gastier-Foster, J., Grody, W.W., Hegde, M., Lyon, E., Spector, E., et al. (2015). Standards and guidelines for the interpretation of sequence variants: A joint consensus recommendation of the American College of Medical Genetics and Genomics and the Association for Molecular Pathology. Genet. Med. 17, 405–424. 10.1038/gim.2015.30.

2. Tavtigian, S.V., Greenblatt, M.S., Harrison, S.M., Nussbaum, R.L., Prabhu, S.A., Boucher, K.M., and Biesecker, L.G.; ClinGen Sequence Variant Interpretation Working Group (2018). Modeling the ACMG/AMP variant classification guidelines as a Bayesian classification framework. Genet. Med. 20, 1054–1060. 10.1038/gim.2017.210.

3. Pejaver, V., Byrne, A.B., Feng, B.-J., Pagel, K.A., Mooney, S.D., Karchin, R., O’Donnell-Luria, A., Harrison, S.M., Tavtigian, S.V., Greenblatt, M.S., et al. (2022). Calibration of computational tools for missense variant pathogenicity classification and ClinGen recommendations for PP3/BP4 criteria. Am. J. Hum. Genet. 109, 2163–2177. 10.1016/j.ajhg.2022.10.013.

4. Bergquist, T., Stenton, S.L., Nadeau, E.A.W., Byrne, A.B., Greenblatt, M.S., Harrison, S.M., Tavtigian, S.V., O’Donnell-Luria, A., Biesecker, L.G., Radivojac, P., et al. (2025). Calibration of additional computational tools expands ClinGen recommendation options for variant classification with PP3/BP4 criteria. Genet. Med. 27, 101402. 10.1016/j.gim.2025.101402.

5. Abderrazzaq, H., Singh, M., Babb, L., Bergquist, T., Brenner, S.E., Pejaver, V., O’Donnell-Luria, A., and Radivojac, P.; ClinGen Computational Working Group; ClinGen Variant Classification Working Group (2026). Calibration of in-frame indel variant effect predictors for clinical variant classification. bioRxiv, 10.64898/2026.04.15.718599.

6. Villani, R.M., McKenzie, M.E., Davidson, A.L., and Spurdle, A.B. (2024). Regional-specific calibration enables application of computational evidence for clinical classification of 5′ cis-regulatory variants in Mendelian disease. Am. J. Hum. Genet. 111, 1301–1315. 10.1016/j.ajhg.2024.05.002.

7. Jaganathan, K., Kyriazopoulou Panagiotopoulou, S., McRae, J.F., Darbandi, S.F., Knowles, D., Li, Y.I., Kosmicki, J.A., Arbelaez, J., Cui, W., Schwartz, G.B., et al. (2019). Predicting splicing from primary sequence with deep learning. Cell 176, 535–548.e24. 10.1016/j.cell.2018.12.015.

8. Walker, L.C., de la Hoya, M., Wiggins, G.A.R., Lindy, A., Vincent, L.M., Parsons, M.T., Canson, D.M., Bis-Brewer, D., Cass, A., Tchourbanov, A., et al. (2023). Using the ACMG/AMP framework to capture evidence related to predicted and observed impact on splicing: Recommendations from the ClinGen SVI Splicing Subgroup. Am. J. Hum. Genet. 110, 1046–1067. 10.1016/j.ajhg.2023.06.002.

9. Hirschi, O.R., Felker, S.A., Rednam, S.P., Vallance, K.L., Parsons, D.W., Roy, A., Cooper, G.M., and Plon, S.E. (2024). Combined bioinformatic and splicing analysis of likely benign intronic and synonymous variants reveals evidence for pathogenicity. Genet. Med. Open 2, 101850. 10.1016/j.gimo.2024.101850.

10. Avsec, Ž., Latysheva, N., Cheng, J., Novati, G., Taylor, K.R., Ward, T., Bycroft, C., Nicolaisen, L., Arvaniti, E., Pan, J., et al. (2026). Advancing regulatory variant effect prediction with AlphaGenome. Nature 649, 1206–1218. 10.1038/s41586-025-10014-0.

11. Cheng, J., Novati, G., Pan, J., Bycroft, C., Žemgulytė, A., Applebaum, T., Pritzel, A., Wong, L.H., Zielinski, M., Sargeant, T., et al. (2023). Accurate proteome-wide missense variant effect prediction with AlphaMissense. Science 381, eadg7492. 10.1126/science.adg7492.

12. Cheng, J., Taylor, K.R., Nicolaisen, L., Pan, J., Bycroft, C., Perino, M., Ward, T., Hawkes, G., Covill, L.E., Weilert, M., et al. (2026). AlphaGenome Atlas: *In silico* mutagenesis of the entire human genome improves prioritization and interpretation of non-coding variants. medRxiv, 10.64898/2026.09.16.26363192.

13. Jaganathan, K., Chen, J., Liu, X., Zhang, Y., Kim, Y., Banerjee, A., Fiziev, P.P., Wilcox, N., Taskiran, I.I., Cai, Y., et al. (2026). A unified framework for quantitative splicing and transcript prediction. bioRxiv, 10.64898/2026.10.08.757673.

14. Cavus, O.A., and Kuskucu, A. (2026). Benchmarking the AlphaGenome Variant Impact score against SpliceAI on post-release ClinVar classifications and experimental splicing assays. bioRxiv, 10.64898/2026.10.11.758362.

15. Quarantani, G., Clarke, J., Thompson, M., Sang, F., Valcárcel, J., and Lehner, B. (2026). OpenSplice: The impact of half a million mutations on the alternative splicing of 600 human exons. bioRxiv, 10.64898/2026.05.22.727141.

16. Tavtigian, S.V., Harrison, S.M., Boucher, K.M., and Biesecker, L.G. (2020). Fitting a naturally scaled point system to the ACMG/AMP variant classification guidelines. Hum. Mutat. 41, 1734–1737. 10.1002/humu.24088.

17. Benegas, G., Eraslan, G., and Song, Y.S. (2025). Benchmarking DNA sequence models for causal regulatory variant prediction in human genetics. bioRxiv, 10.1101/2025.02.11.637758.

18. Chen, S., Francioli, L.C., Goodrich, J.K., Collins, R.L., Kanai, M., Wang, Q., Alföldi, J., Watts, N.A., Vittal, C., Gauthier, L.D., et al. (2024). A genomic mutational constraint map using variation in 76,156 human genomes. Nature 625, 92–100. 10.1038/s41586-023-06045-0.

19. Feng, B.-J. (2017). PERCH: A unified framework for disease gene prioritization. Hum. Mutat. 38, 243–251. 10.1002/humu.23158.

20. Pejaver, V., Urresti, J., Lugo-Martinez, J., Pagel, K.A., Lin, G.N., Nam, H.-J., Mort, M., Cooper, D.N., Sebat, J., Iakoucheva, L.M., et al. (2020). Inferring the molecular and phenotypic impact of amino acid variants with MutPred2. Nat. Commun. 11, 5918. 10.1038/s41467-020-19669-x.

21. Ioannidis, N.M., Rothstein, J.H., Pejaver, V., Middha, S., McDonnell, S.K., Baheti, S., Musolf, A., Li, Q., Holzinger, E., Karyadi, D., et al. (2016). REVEL: An ensemble method for predicting the pathogenicity of rare missense variants. Am. J. Hum. Genet. 99, 877–885. 10.1016/j.ajhg.2016.08.016.

22. Carter, H., Douville, C., Stenson, P.D., Cooper, D.N., and Karchin, R. (2013). Identifying Mendelian disease genes with the Variant Effect Scoring Tool. BMC Genomics 14 (Suppl 3), S3. 10.1186/1471-2164-14-S3-S3.

23. Brandes, N., Goldman, G., Wang, C.H., Ye, C.J., and Ntranos, V. (2023). Genome-wide prediction of disease variant effects with a deep protein language model. Nat. Genet. 55, 1512–1522. 10.1038/s41588-023-01465-0.

24. Wu, Y., Liu, H., Li, R., Sun, S., Weile, J., and Roth, F.P. (2021). Improved pathogenicity prediction for rare human missense variants. Am. J. Hum. Genet. 108, 1891–1906. 10.1016/j.ajhg.2021.08.012.

25. Rentzsch, P., Schubach, M., Shendure, J., and Kircher, M. (2021). CADD-Splice— improving genome-wide variant effect prediction using deep learning-derived splice scores. Genome Med. 13, 31. 10.1186/s13073-021-00835-9.

26. Schubach, M., Maass, T., Nazaretyan, L., Röner, S., and Kircher, M. (2024). CADD v1.7: Using protein language models, regulatory CNNs and other nucleotide-level scores to improve genome-wide variant predictions. Nucleic Acids Res. 52, D1143–D1154. 10.1093/nar/gkad989.

27. Landrum, M.J., Lee, J.M., Benson, M., Brown, G.R., Chao, C., Chitipiralla, S., Gu, B., Hart, J., Hoffman, D., Jang, W., et al. (2018). ClinVar: Improving access to variant interpretations and supporting evidence. Nucleic Acids Res. 46, D1062–D1067. 10.1093/nar/gkx1153.

28. Morales, J., Pujar, S., Loveland, J.E., Astashyn, A., Bennett, R., Berry, A., Cox, E., Davidson, C., Ermolaeva, O., Farrell, C.M., et al. (2022). A joint NCBI and EMBL-EBI transcript set for clinical genomics and research. Nature 604, 310–315. 10.1038/s41586-022-04558-8.

29. Vaz-Drago, R., Custódio, N., and Carmo-Fonseca, M. (2017). Deep intronic mutations and human disease. Hum. Genet. 136, 1093–1111. 10.1007/s00439-017-1809-4.

30. Christmas, M.J., Kaplow, I.M., Genereux, D.P., Dong, M.X., Hughes, G.M., Li, X., Sullivan, P.F., Hindle, A.G., Andrews, G., Armstrong, J.C., et al. (2023). Evolutionary constraint and innovation across hundreds of placental mammals. Science 380, eabn3943. 10.1126/science.abn3943.

31. You, N., Liu, C., Gu, Y., Wang, R., Jia, H., Zhang, T., Jiang, S., Shi, J., Chen, M., Guan, M.-X., et al. (2024). SpliceTransformer predicts tissue-specific splicing linked to human diseases. Nat. Commun. 15, 9129. 10.1038/s41467-024-53088-6.

32. Zeiberg, D., Jain, S., and Radivojac, P. (2020). Fast nonparametric estimation of class proportions in the positive-unlabeled classification setting. Proc. AAAI Conf. Artif. Intell. 34, 6729–6736. 10.1609/aaai.v34i04.6151.

33. Truty, R., Ouyang, K., Rojahn, S., Garcia, S., Colavin, A., Hamlington, B., Freivogel, M., Nussbaum, R.L., Nykamp, K., and Aradhya, S. (2021). Spectrum of splicing variants in disease genes and the ability of RNA analysis to reduce uncertainty in clinical interpretation. Am. J. Hum. Genet. 108, 696–708. 10.1016/j.ajhg.2021.03.006.

