## Supplemental Methods and Results for "Calibration of AlphaGenome Variant Impact and SpliceAI2 scores as PP3/BP4 evidence for missense, non-coding, and synonymous variants"

Abbreviations (AVI, PP3, BP4, P/LP, B/LB), the analysis sets (primary set, published sets), and the training-related exclusion and rarity rule are as defined in the main text. AVI scores are given on the PHRED scale of the AVI release unless stated otherwise.

### Local posterior probability method

We used the local posterior probability method of Pejaver et al. (2022) and Bergquist et al. (2025). For each candidate threshold, the proportion of pathogenic variants is estimated in the smallest score window around it that contains at least 100 labeled variants and at least 3% of an unlabeled reference set of population variants. Benign variants are weighted so that pathogenic variants make up 4.41% of the weighted total, the prior probability of pathogenicity. The weighted proportion of pathogenic variants in a window is then a local posterior probability. Each posterior probability corresponds to exactly one local positive likelihood ratio (LR). For an evidence level worth $n$ points, the required LR is $c^{n/8}$, where $c$ represents the LR for very strong evidence. Following Pejaver et al. (2022), $c$ was set to 1,124, the value they reported for a prior probability of pathogenicity of 4.41% for rare missense variants. Because $c$ is fixed, the prior used for the weights does not change the thresholds. The threshold of a pathogenic evidence level is the lowest score from which the local LR meets the requirement at every higher candidate score; benign thresholds are defined in the same way from the low end of the score range.

The window was widened in fixed steps. The step was the largest power of ten that divides the range between the 0.5th and 99.5th percentiles of the reference scores into at least 1,000 steps. Scores were placed on a grid of one tenth of this step (AVI raw scores: step 0.001, grid 0.0001; SpliceAI2: step 0.0001, grid 0.00001). For the other missense tools, we used the window steps of Pejaver et al. (2022): 0.01 for BayesDel, 0.1 for CADD, and 0.001 for the remaining tools. For ESM1b, which they did not calibrate, the step was 0.01. Scores were used at their published precision, except that MutPred2 and VARITY_R scores were rounded to 0.0001. We also recalculated the requirements for assumed priors of 0.5%, 1%, 2%, 4.41%, and 10% (Tables S9 and S10), with $c = (9(1 - X)/X)^{4/3}$ for prior $X$, following Tavtigian et al. (2018).

The ACMG/AMP guidelines do not define an evidence level corresponding to 3 points. In the points-based framework (Tavtigian et al., 2020), this level falls between moderate and strong evidence; we refer to it as +3, as in Bergquist et al. (2025), and as −3 for benign evidence.

Reported thresholds are the one-sided 95% bootstrap bounds, which are conservative estimates of the scores required to reach each evidence level; an evidence level was considered reached only if it was reached in at least 95% of bootstrap replicates. The bound is the 95th percentile of the bootstrap thresholds in the stricter direction (higher scores for pathogenic and lower scores for benign levels), with replicates in which the level was not reached counted as the strictest. We used 10,000 bootstrap replicates that resampled variants for the main missense analysis and 2,000 replicates that resampled genes for the non-coding and synonymous classes. As in the original code of the method, the weight of benign variants was calculated once from all labeled variants and kept the same in every replicate; recalculating it in each gene-resampled replicate would have added one evidence level to the main non-coding and synonymous analysis (SpliceAI2 benign supporting evidence in deep introns, threshold 0.00). The other thresholds would have changed by at most 0.01 for SpliceAI2 and by at most 0.14 on the PHRED scale for AVI, except two AVI thresholds: deep intronic pathogenic supporting (24.54 instead of 25.11) and benign strong at near-canonical positions (2.28 instead of 9.67). An evidence level was not counted if its threshold was reached by no more than one labeled variant (for a pathogenic level, a pathogenic variant scoring at or above the threshold; for a benign level, a benign variant scoring at or below it). Such thresholds lie at the edge of the available data and are flagged in the supplemental tables.

To check our implementation, we recalculated the point-estimate thresholds published by Pejaver et al. (2022) for 13 tools and eight levels (104 thresholds), with scores placed on an exact grid at their published precision. Neither calibration reached 45 of these levels. Of the 59 levels that were reached, 51 thresholds were identical at the published precision. Five differed by at most 0.0021: four because the published input scores have limited precision (PrimateAI and MutPred2), and one (EA1.0 benign supporting, 0.2865 vs. 0.286) because our value lies exactly halfway between two values at the published precision. Three differed because a score lay exactly on a window edge (ours vs. published: FATHMM benign supporting 2.23 vs. 2.20; phyloP benign moderate 0.385 vs. 0.323; VEST4 benign strong 0.072 vs. 0.073).

### Scores

AVI scores were taken from the developers' genome-wide file of single-nucleotide variants (Cheng et al., 2026). Thresholds were calculated on the raw scores and converted to the PHRED scale of the AVI release. PHRED increases with the raw score. Each pathogenic threshold was converted to the lowest PHRED value among variants with the smallest raw score at or above it, and each benign threshold to the highest PHRED value among variants with the largest raw score at or below it, so that the same variants pass the threshold on both scales. PHRED thresholds are given to two decimals, rounded up for pathogenic and down for benign thresholds; values below 1, where many raw scores map to nearly the same PHRED value, are given unrounded. Raw AVI thresholds for the non-coding and synonymous classes are given in Table S9.

SpliceAI2 scores were taken from the developers' precomputed files (Jaganathan et al., 2026). We used the summary score, the largest of the four predicted changes in splice-site use (acceptor or donor gain or loss), and, for variants with scores on both strands, the higher value.

We ran SpliceAI v1.3.1 with a maximum distance of ±4,999 nucleotides and without masking. We took the largest of its four predicted changes in splice-site use, over all genes if the variant lay in several. Precomputed SpliceAI scores (50-nucleotide window) were obtained from the Ensembl VEP REST API, and we took the largest predicted change over all transcripts. For SpliceTransformer, we used its overall splicing score with its own GENCODE v38 annotation, taking the largest value over the strands of overlapping genes. phyloP was taken from the Cactus

241-mammal alignment at the variant position, and CADD v1.7 as PHRED scores. Variants without a score received no evidence.

### Reference set

For the non-coding and synonymous classes, the reference set consisted of gnomAD v4.1 genome variants of the same classes in genes with at least one pathogenic or likely pathogenic classification in ClinVar, on chromosomes not used for AVI training, with an allele frequency below 0.01. Variants had to pass the gnomAD filters and lie outside low-complexity and segmental-duplication regions. Among carriers, the median depth had to be at least 30 (at least 10 for an allele count of 3 or more) and the median genotype quality at least 20. Variants present in ClinVar were excluded. In the main analysis, the reference set was further restricted to rare variants (filtering allele frequency below 0.001; where it was missing, group-maximum allele frequency below 0.001; where that was also missing, global allele frequency below 0.001), as were the labeled variants under the rarity rule; the analysis without the rarity rule used all variants with allele frequency below 0.01. We kept all variants within 200 nucleotides of an exon and a random 5% of variants farther away. Deep intronic variants 101–200 nucleotides from an exon were then subsampled to the same 5% density, so that each class has one sampling density. Each class was limited to 200,000 variants. For missense variants, the reference set was the published gnomAD v2 set.

### Population variants as controls

Because population controls may include undiscovered pathogenic variants, the estimated likelihood ratios are biased toward 1 and cannot exceed the reciprocal of the pathogenic fraction in the control set. The controls were a random sample of 30,000 population variants of the class from the reference set (all of them where fewer were available); the score windows were still sized on the full reference set. We report the uncorrected results, which assume that the controls contain no pathogenic variants. We considered a pathogenic evidence level assessable only when its required likelihood ratio was no greater than half of this maximum, $1/\alpha$, where $\alpha$ is the pathogenic fraction among the controls. This required a pathogenic fraction below 20.8% for supporting, 8.6% for moderate, 3.6% for +3, and 1.5% for strong evidence. Where the estimated upper limit of the pathogenic fraction (Table 2) or, for missense variants, the published prior probability of 4.41% exceeded these limits, levels that were not reached were reported as not assessable rather than as not reached. The rule affected only the +3 and strong levels, for missense variants, near-canonical splice positions, and the exonic splice region. For the polypyrimidine tract, where the estimated upper limit was approximately 2%, it applied only to strong evidence.

For the correction, each required likelihood ratio $L$ (a minimum for pathogenic and a maximum for benign evidence) was replaced by $L/(1 - \alpha + \alpha L)$, where $\alpha$ is the assumed pathogenic fraction among the controls; a corrected pathogenic level was assessable only if this requirement was at most $0.5/\alpha$. With $\alpha = 4.41\%$, the published prior for missense variants, the +3 level becomes assessable, whereas strong evidence does not. Corrected analyses assuming pathogenic fractions of 0.5%–10% are provided for the non-coding and synonymous classes in Table S1; missense variants were corrected using 4.41% (Table S4b).

### Classifications made before SpliceAI

For each variant class, we divided the calibration set into two groups: variants that already had the same classification (pathogenic or likely pathogenic, or benign or likely benign) with at least one review star in the ClinVar release of 2 December 2018, and all remaining variants, most of which were classified or reclassified later. Some variants in the latter group already had classifications in 2018 but had no review star. Within each group, we calculated the likelihood ratios of the pathogenic and of the benign intervals of supporting evidence or higher under the main thresholds (Table S16c), and the area under the receiver operating characteristic curve (AUC) of SpliceAI2 and AVI. We then estimated how the difference in AUC between SpliceAI2 and AVI changed between the two groups, using 300 gene-cluster bootstrap replicates (Table S16d).

Because the earlier classifications were also part of the calibration set, this comparison was not independent of the thresholds. It also tested only one combined interval per direction and class. We therefore also repeated the calibration using only the later classifications, with the same pipeline and reference set, and evaluated every level reached against the earlier classifications with the verdict rules of the validation (Tables S16a and S16b). The two groups contained no common variants but shared some genes and genomic positions. The variant classes and the gene list were defined from all classifications, so this analysis tests only the thresholds, not the class definitions or the gene list.

### Robustness analyses

For missense variants, we repeated the AVI calibration in the following sets (Table S2c): with only one of the two restrictions (training-related exclusion or rarity rule); with the training-related exclusion based only on gnomAD variants that passed quality filters; on the 16 chromosomes not used for AVI training; on the eight chromosomes used neither for training nor for selecting the final model (3, 6, 9, 12, 16, 18, 19, and 21); on variants with an AlphaMissense score; on variants scored by every tool, in the primary and in the published sets; and with rarity defined from gnomAD exomes and genomes together. The AVI developers trained AVI on chromosomes 1, 4, 7, 8, 10, 13, and 15 and selected the final model (checkpoint selection) with data on chromosomes 2, 5, 11, 14, 17, 20, 22, and X, including data for four genes (*BRCA1*, *ATM*, *RAD51C*, and *DDX3X*) whose variants were excluded from the primary set (Cheng et al., 2026). We also removed random variants, matched in number to each restriction and to both, five times each. Finally, we repeated the calibration with a gene-cluster bootstrap, on the PHRED scale, with half and double the window step, and with a score grid of 0.00001 instead of 0.0001. Each analysis used 2,000 bootstrap replicates (1,000 with the finer grid).

For the non-coding and synonymous classes, the analyses in Table S1 that kept the class definitions were the main analysis; without the training-related exclusion; without one-star likely benign classifications; pathogenic against benign classifications only (without likely pathogenic and likely benign); non-training chromosomes only; a reference window of 1% instead of 3%; a second random seed; and without the rarity rule (all variants with allele frequency below 0.01). The other analyses, which changed the class definitions, the bootstrap unit, or the comparison group, were without donor position +5; near-canonical positions and polypyrimidine tract pooled; variants resampled instead of genes; boundaries of 200 and 500 nucleotides between proximal and deep introns, each also with population variants as controls; and population variants as controls, uncorrected and corrected for an assumed pathogenic fraction. A random-removal analysis removed the same share of variants as the corresponding restriction, but at random; it was run three times for each of the two restrictions (Tables S9 and S10). The 13 robustness analyses with unchanged class definitions are the seven analyses from “without the training-related exclusion” to “without the rarity rule” and the six random-removal runs. Finally, we repeated the main calibration with both pathogenic and benign ClinVar variants restricted to those absent from gnomAD v4.1 genomes, to assess whether differences in allele frequency between these groups affected the thresholds (Table S11).

### Comparison of tools

We evaluated further tools on a fixed subset of 326,618 variants (Table S18) and calibrated the original SpliceAI with the same pipeline (Table S19b). The subset was chosen from labels and variant classes only, without using any score; labeled variants were drawn from those with allele frequency below 0.01 after the training-related exclusion, before the rarity rule was applied. It contained all pathogenic variants of the calibration and validation sets, up to 20,000 benign variants per class in each set, and up to about 20,000 reference variants per class. Separately, AVI and SpliceAI2 were compared on all variants scored by both tools using the AUC and sensitivity at 99% specificity, with either ClinVar benign variants or population variants as the negative class; negative sets larger than 20,000 variants were randomly reduced to 20,000 (Table S7). Differences were computed from unrounded values and assessed using 2,000 gene-cluster bootstrap replicates. These comparisons were descriptive, without adjustment for multiple testing.

To assess incremental value in the non-coding and synonymous classes, we used 5-fold cross-validation with folds grouped by gene. Logistic regression models on rank-transformed AVI and SpliceAI2 scores were fitted on four folds and tested on the fifth. We measured the gain in AUC when one score was added to the other, averaging the AUCs over the test folds. Negative sets larger than 20,000 variants were randomly reduced to 20,000. For missense variants, AVI was combined with AlphaMissense or REVEL in a logistic regression model on the raw scores, fitted on ClinVar 2019 variants of the primary set and evaluated on ClinVar 2020 variants absent from the 2019 set. Intervals are from 2,000 gene-cluster bootstrap replicates (Table S6a). We also calculated Spearman rank correlations of AVI with AlphaMissense, CADD v1.7, and REVEL (Table S5).

To explore the contribution of AVI beyond splicing prediction, we counted variants with an AVI score at or above the pathogenic supporting threshold of their class and a SpliceAI2 score below 0.05 (and, more strictly, at most 0.01), overall and split into classifications made before SpliceAI and later ones (Table S6b).

### Benign classifications, population controls, and prior probability

For proximal and deep intronic variants, we calculated the shares of ClinVar pathogenic, ClinVar benign, and population variants that scored at or above the pathogenic supporting threshold obtained with population controls (Table S14, which also gives the higher levels). We calculated these shares within bins of distance from the exon (21–50 and 51–100 nucleotides in proximal introns; 101–200, 201–500, 501–2,000, and more than 2,000 nucleotides in deep introns), and again with population variants restricted to genes with labeled variants. For deep intronic variants, we also measured how concentrated the variants were in genes and recalculated AUCs after excluding the five and ten genes with the most pathogenic variants (Table S15).

For the prior probability of pathogenicity, we calculated three measures. The floor was the share of rare population variants already classified as pathogenic or likely pathogenic. The ceiling was an upper confidence limit, calculated separately for AVI and SpliceAI2 (Table 2). At each of the nine deciles of the pathogenic scores, we divided the upper 97.5% Jeffreys limit of the share of population variants at or above that score by the lower 2.5% Jeffreys limit of the share of pathogenic variants at or above it. The ceiling was the smallest of these nine ratios. The ceiling assumes that undiscovered pathogenic variants score like known ones. The third measure was a mixture-proportion estimate with DistCurve (Zeiberg et al., 2020). Its input was a logistic-regression combination of five rank-transformed scores other than AVI and SpliceAI2 (SpliceAI with the ±4,999-nucleotide window and as precomputed scores, SpliceTransformer, CADD v1.7, and phyloP). Each variant's combined score came from a model fitted without that variant (cross-validation), on the fixed subset. Uncertainty came from 100 bootstrap replicates.

### Performance of AVI on its training chromosomes (missense variants)

AVI was trained on allele-frequency labels from gnomAD on chromosomes 1, 4, 7, 8, 10, 13, and 15. To test whether it behaves differently for variants it may have seen in training, we used the missense variants of ClinVar 2019 and 2020 before the training-related exclusion and the rarity rule, without the four genes used for checkpoint selection. Each variant was scored as receiving pathogenic evidence (score at or above the tool's pathogenic supporting threshold in the published sets; Table S2a), benign evidence (at or below the benign supporting threshold), or neither. For each group of variants, we calculated the separation: half the difference between the natural log likelihood ratios of pathogenic and of benign evidence. On the training chromosomes, we took the separation for variants present in gnomAD v4.1 genomes minus that for absent variants, and subtracted the same difference on the eight chromosomes used neither for training nor for selecting the final model (3, 6, 9, 12, 16, 18, 19, and 21). Intervals are from a gene-cluster bootstrap (2,000 replicates); the two-sided $p$ value is from 500 random splits of these 15 chromosomes into sets of seven and eight. The same analysis was run for three tools whose training did not separate these chromosomes from the others (AlphaMissense, BayesDel, and CADD v1.6; Table S22).

The difference had the same sign and a similar size for all four tools and lay within the range seen for random sets of seven chromosomes ($p$ = 0.05–0.14), so it is not specific to AVI and gives no indication that AVI performs better on variants it may have seen in training.

**Table S22. Performance of AVI and three other missense tools on the AVI training chromosomes.** Separation is half the difference between the natural log likelihood ratios of a tool's pathogenic and benign supporting evidence (thresholds from the published sets, Table S2a). The difference in differences is the separation for variants present in gnomAD v4.1 genomes minus that for absent variants on the seven AVI training chromosomes, minus the same difference on the eight chromosomes used neither for training nor for selecting the final model. A positive value would indicate better performance on training-chromosome variants seen in gnomAD. Variants: ClinVar 2019 and 2020 missense variants before the training-related exclusion and the rarity rule, without the four checkpoint genes. 95% intervals from 2,000 gene-cluster bootstrap replicates; two-sided *p* from 500 random splits of the 15 chromosomes into sets of seven and eight.

| Tool | Difference in differences | 95% interval | Permutation *p* |
|---|---|---|---|
| AVI | −0.38 | −0.80 to 0.06 | 0.14 |
| AlphaMissense | −0.49 | −0.96 to −0.03 | 0.13 |
| BayesDel | −0.44 | −0.85 to −0.02 | 0.05 |
| CADD v1.6 | −0.42 | −0.84 to 0.00 | 0.06 |

### Supplemental results

#### Comparator tools for missense variants

In the published sets, the thresholds of the seven tools recommended by ClinGen closely reproduced their published values (Pejaver et al., 2022; Bergquist et al., 2025), except that ESM1b did not reach strong evidence (Table S2a). In the primary set, their pathogenic thresholds were slightly stricter, and AlphaMissense, MutPred2, and VARITY_R reached +3 rather than strong evidence. The benign strong levels of REVEL and MutPred2 and the benign −3 level of ESM1b were reached by at most one labeled benign variant and were therefore not counted (Table S2a; Figure 2A).

#### Higher evidence levels in proximal introns and at other synonymous sites

Other synonymous sites had fewer than 300 pathogenic variants, so levels above moderate could not be estimated. The calibration nevertheless produced thresholds for +3 evidence (AVI 24.26; SpliceAI2 0.41) and strong evidence (SpliceAI2 0.66; Tables S9 and S10). AVI also gave a strong threshold of 49.53, but this is the highest AVI score in the class, held by a single benign variant, so no pathogenic variant reached it. Among the validation variants at these sites (classified in the ClinVar release of 28 September 2026 but not in the 2025 release; 37 pathogenic and 62,625 benign variants), scores in these ranges were much more common among pathogenic variants. In the AVI +3 range there were 15 pathogenic and 59 benign variants (likelihood ratio 430). In the SpliceAI2 +3 range there were 11 and 29 (642), and in the SpliceAI2 strong range 10 and 5 (3,390). These counts were obtained by applying the thresholds to the validation set as in Table S3; they are not shown in the tables, because these levels were not estimable. These classifications may have used splicing predictions. Proximal introns also had fewer than 300 pathogenic variants (135). There, AVI gave a +3 threshold of 27.07 (Table S9), likewise not estimable, and SpliceAI2 gave no +3 threshold (Table S10); neither tool gave a strong threshold.

#### Variation of thresholds across robustness analyses

Even in the 13 analyses with unchanged class definitions, the SpliceAI2 supporting threshold in the polypyrimidine tract ranged from 0.19 to 0.30, with the upper value matching the moderate threshold in the main analysis, and SpliceAI2 reached strong evidence at near-canonical positions in eight of the 13 (Tables S1 and S10).

#### Gene concentration of high-scoring deep intronic benign variants

The 974 deep intronic benign variants with AVI scores at or above the population-control supporting threshold (14.43) were distributed across 208 genes and were not more concentrated in a small number of genes than deep intronic benign variants overall. The ten most frequently represented genes accounted for 34% of these high-scoring variants, compared with 37% of all deep intronic benign variants. In addition, excluding genes with the highest numbers of pathogenic variants changed the AUC of either tool by up to 0.03 (Table S15).

#### Likelihood ratio for splice disruption

At near-canonical positions, stronger pathogenic evidence also meant more splice disruption: the likelihood ratio for disruption rose from 2.7 in the supporting interval to 7.9 in the strong interval for AVI, and from 2.0 to 8.1 (supporting to +3, its highest level) for SpliceAI2 (Table S21).

### Notes on Tables S9 and S10

*Analysis:* main analysis or robustness analysis (names as in Table S1; the analysis with variants resampled is in Table S1 only, and random-removal replicates are listed only here).

*Variant class:* as defined in the main text.

*Requirements and benign group:* "Main requirements, ClinVar benign", the requirements used in the main text ($c$ = 1,124) with ClinVar benign classifications; "Main requirements, population controls, uncorrected" and "Main requirements, population controls, corrected for $\alpha$", population variants as controls, uncorrected or corrected for an assumed pathogenic fraction $\alpha$; "Requirements for a prior of $X$, ClinVar benign", requirements recalculated for an assumed prior probability of pathogenicity $X$ (0.5%, 1%, 2%, 4.41%, or 10%) with $c = (9(1 - X)/X)^{4/3}$, following Tavtigian et al. (2018); this gives 1,131 at 4.41%, whereas Pejaver et al. (2022) used 1,124.

*c (LR for very strong evidence):* likelihood ratio for very strong evidence underlying the requirements; blank in the boundary analyses.

*Direction, Points:* direction (pathogenic, PP3; benign, BP4) and evidence level in points (1, supporting; 2, moderate; 3, +3 for pathogenic or −3 for benign evidence; 4, strong; 8, very strong).

*Point estimate, Threshold:* point estimate of the threshold and its one-sided 95% bootstrap bound; Table S9 gives both as raw AVI scores and as PHRED equivalents.

*Failed bootstrap replicates (share):* share of bootstrap replicates in which the level was not reached.

*Estimable (sample-size rule):* whether the level could be estimated under the sample-size rule.

*Assessable:* population-control analyses only (blank in the others); yes if the level was assessable at the assumed fraction, no if not; always yes for benign levels; blank for uncorrected pathogenic levels (see the next column) and in the boundary analyses.

*Largest pathogenic fraction among controls at which assessable:* uncorrected population-control analysis only; the largest pathogenic fraction among the controls at which the pathogenic level would be assessable (0.5 divided by the required likelihood ratio); blank in the boundary analyses.

*P/LP, B/LB, Reference variants:* numbers of pathogenic, benign, and reference variants (B/LB is 0 in population-control rows, where the controls were a random sample of up to 30,000 of the reference variants).

*Bootstrap replicates, Random seed, Reference window (share of reference set):* number of bootstrap replicates, random seed, and reference window; blank in the boundary analyses.

*Not counted:* evaluated only for estimable levels that were reached; yes if at most one labeled variant of the level's direction in the class reaches the threshold, and such levels are not counted; no otherwise.

### Supplemental references

Bergquist, T., Stenton, S.L., Nadeau, E.A.W., Byrne, A.B., Greenblatt, M.S., Harrison, S.M., Tavtigian, S.V., O'Donnell-Luria, A., Biesecker, L.G., Radivojac, P., et al. (2025). Calibration of additional computational tools expands ClinGen recommendation options for variant classification with PP3/BP4 criteria. Genet. Med. 27, 101402. 10.1016/j.gim.2025.101402.

Cheng, J., Taylor, K.R., Nicolaisen, L., Pan, J., Bycroft, C., Perino, M., Ward, T., Hawkes, G., Covill, L.E., Weilert, M., et al. (2026). AlphaGenome Atlas: *In silico* mutagenesis of the entire human genome improves prioritization and interpretation of non-coding variants. medRxiv, 10.64898/2026.09.16.26363192.

Jaganathan, K., Chen, J., Liu, X., Zhang, Y., Kim, Y., Banerjee, A., Fiziev, P.P., Wilcox, N., Taskiran, I.I., Cai, Y., et al. (2026). A unified framework for quantitative splicing and transcript prediction. bioRxiv, 10.64898/2026.10.08.757673.

Pejaver, V., Byrne, A.B., Feng, B.-J., Pagel, K.A., Mooney, S.D., Karchin, R., O'Donnell-Luria, A., Harrison, S.M., Tavtigian, S.V., Greenblatt, M.S., et al. (2022). Calibration of computational tools for missense variant pathogenicity classification and ClinGen recommendations for PP3/BP4 criteria. Am. J. Hum. Genet. 109, 2163–2177. 10.1016/j.ajhg.2022.10.013.

Tavtigian, S.V., Greenblatt, M.S., Harrison, S.M., Nussbaum, R.L., Prabhu, S.A., Boucher, K.M., and Biesecker, L.G.; ClinGen Sequence Variant Interpretation Working Group (2018). Modeling the ACMG/AMP variant classification guidelines as a Bayesian classification framework. Genet. Med. 20, 1054–1060. 10.1038/gim.2017.210.

Tavtigian, S.V., Harrison, S.M., Boucher, K.M., and Biesecker, L.G. (2020). Fitting a naturally scaled point system to the ACMG/AMP variant classification guidelines. Hum. Mutat. 41, 1734–1737. 10.1002/humu.24088.

Walker, L.C., de la Hoya, M., Wiggins, G.A.R., Lindy, A., Vincent, L.M., Parsons, M.T., Canson, D.M., Bis-Brewer, D., Cass, A., Tchourbanov, A., et al. (2023). Using the ACMG/AMP framework to capture evidence related to predicted and observed impact on splicing: Recommendations from the ClinGen SVI Splicing Subgroup. Am. J. Hum. Genet. 110, 1046–1067. 10.1016/j.ajhg.2023.06.002.

Zeiberg, D., Jain, S., and Radivojac, P. (2020). Fast nonparametric estimation of class proportions in the positive-unlabeled classification setting. Proc. AAAI Conf. Artif. Intell. 34, 6729–6736. 10.1609/aaai.v34i04.6151.
